# Non-coding RNAs drive transcriptional network rewiring underlying thermal dimorphism in a human pathogenic fungus

**DOI:** 10.64898/2026.09.29.755541

**Authors:** Xueyan Hu, Juan Wang, Ence Yang

## Abstract

As fungal pathogens pose mounting global challenges, decoding their complex adaptive traits remains a paramount academic priority. While previous studies have provided foundational insights into protein-coding variations and orphan genes, fully understanding such traits requires exploring upstream non-coding regulatory networks. Here, we present a high-resolution, rRNA-depleted temporal transcriptomic framework across reciprocal dimorphic transitions to decode how non-coding network rewiring involving in fungal adaption. Integrating temporal profiling with comparative genomics reveals that early ATP-related metabolic reprogramming and melanin biosynthesis direct morphogenesis via species-specific motifs. Further temporal network modeling uncovers a functional stratification where long non-coding RNAs orchestrate early stress responses, whereas circular RNAs drive morphological conversion. Functional validation confirms that hub non-coding transcripts mediate stress adaptation, with cross-species perturbation in the closest non-pathogen species altering phenotypic outcomes. Together, these findings demonstrate how system-level non-coding dynamics underlie the evolutionary innovation of thermal dimorphism, establishing a robust paradigm for understanding specialized fungal pathogenicity and evolution adaption.

## Introduction

Human fungal pathogens pose a severe burden on global human health, causing millions of life-threatening systemic and superficial infections annually^1^. Compared with non-pathogenic counterparts, pathogenic fungi require profound physiological and morphological adaptations against multifaceted host microenvironments and immune defenses^2^. However, protein-coding variations and orphan genes alone are frequently insufficient to fully account for such multifaceted adaptive traits^3,4^, a limitation starkly exemplified by the unique dimorphic fungus *Talaromyces marneffei* within Eurotiales, where the limited repertoire of hundreds of orphan genes relative to non-pathogenic relatives contrasts sharply with complex pathogenesis and morphological transitions^5–7^. Overcoming this explanatory gap necessitates complementing isolated gene turnover analyses with gene regulatory network reconstruction, recognizing that pathogen-specific phenotypes are shaped significantly by modular network rewiring^8^. Increasing evidence indicates that non-coding elements and their associated RNA networks act as central hubs driving these rewiring events, modulating *cis*-regulatory landscapes and binding scaffolds to confer specialized host adaptability^9–12^. While recent microevolutionary investigations have demonstrated the essential roles of several non-coding RNAs in fungal phenotypic plasticity and environmental adaptation^13–15^, building upon these foundations requires a broader macroevolutionary perspective to decipher how non-coding networks drive the evolutionary origins of specialized fungal pathogenicity.

As the unique thermally dimorphic pathogen among over one thousand species within the order Eurotiales, *T. marneffei* provides a powerful comparative framework for dissecting evolutionary and pathogenic transitions^4^. Its ability to undergo temperature-dependent morphological transitions is a hallmark of pathogenicity^16^. At ambient temperatures (25°C), *T. marneffei* exists as a filamentous, saprophytic form, reproducing through conidia, whereas exposure to mammalian body temperature (37°C) induces a tightly regulated switch toward a yeast-like parasitic morphology, highlighting exceptional phenotypic plasticity^17^. Previous poly(A)-based temporal transcriptomics, which initially focusing primarily on protein-coding trajectories, have provided valuable expression snapshots of downstream phenotypic outcomes such as cell wall remodeling and metabolic reprogramming^18^. Moreover, specific molecular investigations established the regulatory functions of specific non-coding transcripts in morphogenesis, as exemplified by the yeast-enriched transcript circDS-1^14^. Advancing beyond these insights highlights the necessity of systematically integrating comprehensive coding and non-coding profiling to delineate the upstream non-coding regulatory architectures governing thermal transitions, establishing a framework to elucidate how global non-coding network rewiring drives the evolutionary innovation of thermal dimorphism of *T. marneffei*. Here, we generated a high-quality rRNA-depleted RNA-seq dataset capturing the temporal dynamics of the dimorphic transition in *T. marneffei* across fifteen time points spanning zero to 108 hours in both the mycelium-to-yeast and yeast-to-mycelium directions. Time-series differential expression and phylogenetic comparative analyses delineated precise pathway transitions unique to *T. marneffei*, revealing that early ATP-related metabolic reprogramming and melanin biosynthesis serve as essential determinants for morphogenesis driven by species-specific stress-related promoters and motifs. Trans-acting network analyses demonstrated that lncRNAs and circRNAs orchestrate distinct temporal layers of regulatory rewiring, with lncRNAs operating during the early preparatory phase and circRNAs driving the subsequent late morphological conversion phase. Functional validation experiments further established that three hub non-coding RNAs within the early regulatory network exert stress-related functions during the dimorphic transition in *T. marneffei*, with one prominent stress-protection lncRNA displaying altered phenotypic outcomes upon perturbation in the most closely related non-pathogenic species *T. stipitatus*. Together, these findings demonstrate how system-level molecular dynamics uncover the core regulatory architecture driving environmental adaptation in *T. marneffei*, establishing a definitive paradigm for how non-coding network rewiring underlies the evolutionary innovation of fungal thermal dimorphism.

## Results

### Comprehensive whole-transcriptome time-series profiling reveals dynamic landscapes during *T. marneffei* dimorphic transition

To decode the non-coding RNA-mediated transcriptomic rewiring underlying the *T. marneffei* dimorphic transition, we employed the established rProbe-seq methodology to construct ribosomal RNA-depleted transcriptomic datasets across multiple temporal points (0, 0.5, 1, 2, 3, 6, 9, 12, 24, 36, 48, 60, 72, 84, 108 hours) during both yeast-to-mycelium (Y2M) and mycelium-to-yeast (M2Y) transitions^19^ (Figure 1A and Figure S1). Each sample generated an average of 65 million clean reads, ensuring robust transcript detection and quantification. High correlation coefficients between biological replicates (*R* > 0.9) and temporally adjacent samples demonstrated excellent sequencing quality and experimental reproducibility (Figure 1B).

**Figure 1.**
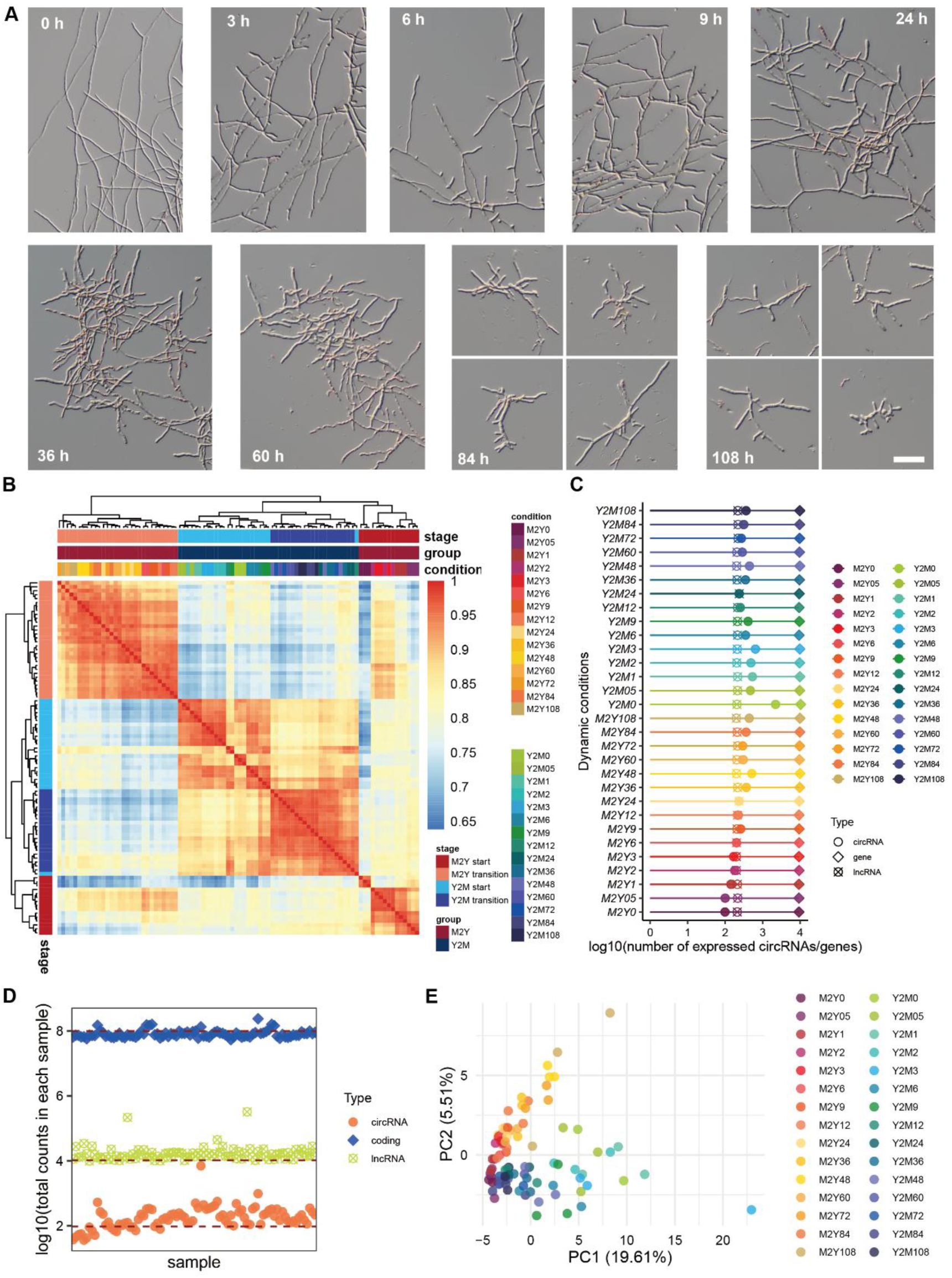
Times series rRNA-depletion RNA-seq of dimorphic transition of *T. marneffei*. (A) Morphological changes of *T. marneffei* during the M2Y time series cultivation. Scale bar = 50 μm. (B) Cluster heatmap of transcriptomic samples. Color legend indicates Spearman correlation coefficients between samples. The x-axis and y-axis represent the time-series transcriptome samples. (C) Numbers of protein-coding genes, lncRNAs, and circRNAs expressed in each transcriptomic sample of the time series. (D) Percentage counts of protein-coding genes, lncRNAs, and circRNAs expressed in each sample across time series. (E) Principal component analysis (PCA) of circRNA expression matrices obtained from the time-series data.

**Figure S1.**
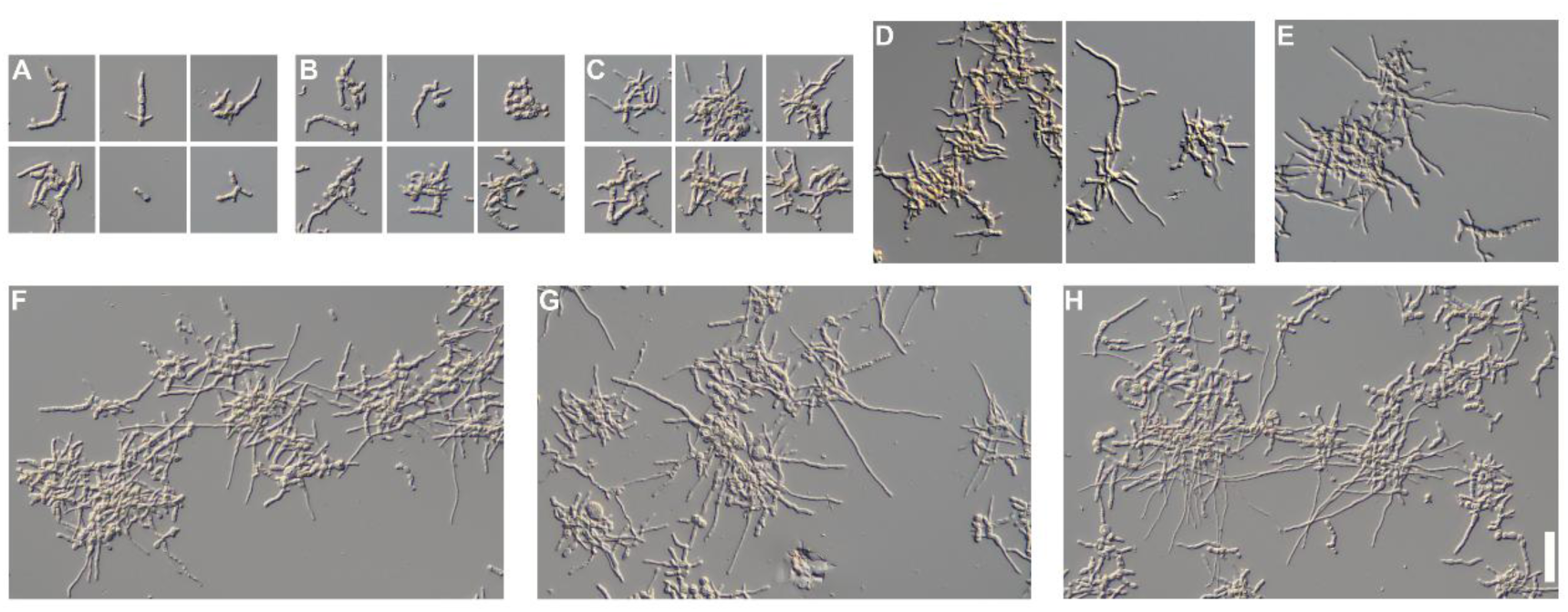
Morphological dynamics during yeast-to-mycelium transition in *T. marneffei*. (A-I) Microscopic images showing morphological changes during yeast-to-mycelium (Y2M) transition at specific time points: 0 h, 3 h, 6 h, 9 h, 24 h, 36 h, 60 h, 84 h, and 108 h. Scale bar = 50 μm.

**Figure S2.**
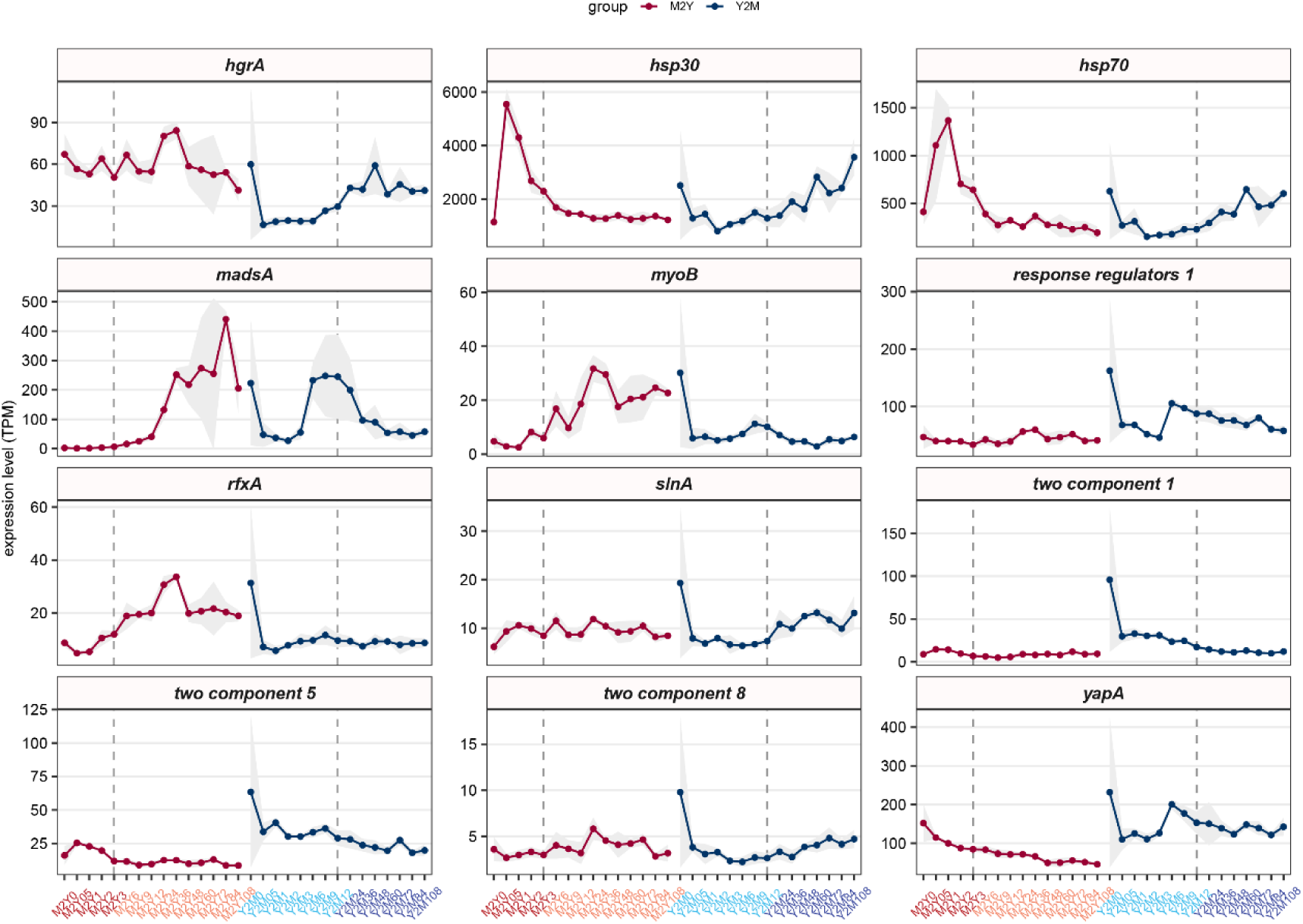
Expression patterns of previously reported dimorphism-associated genes. Line plot showing TPM-based expression dynamics of reported dimorphism-associated genes. The x-axis represents time points, while the y-axis indicates mean TPM values across biological replicates (mean ± SD shown in gray shading).

To validate the biological representativeness of our morphological cultivation system, we examined the expression profiles of previously characterized dimorphic transition regulators within our temporal dataset (Figure S2). These established markers exhibited characteristic differential expression patterns corresponding to their known functional phases. Specifically, *hgrA*, encoding a C2H2 zinc finger transcription factor that promotes hyphal development^20^, demonstrated progressive upregulation during late-stage Y2M transition. The heat shock proteins *hsp30* and *hsp70* showed rapid temperature-induced expression elevation during early M2Y phases^21^, while *myoB*^22^, which regulates chitin accumulation, exhibited reciprocal upregulation and downregulation patterns during M2Y and Y2M transitions, respectively, reflecting the cell wall remodeling processes characteristic of mycelial and yeast cell morphotypes. These established markers exhibited characteristic differential expression patterns corresponding to their known functional phases, confirming the biological validity of our newly established liquid cultivation system.

Through comprehensive transcriptomic analysis, we identified 10,347 protein-coding genes, 6,930 circRNAs, and 240 intergenic lncRNAs that exhibited detectable expression in at least one temporal point (Figure 1C). Intergenic lncRNAs exhibited greater length heterogeneity (range: 202 nt to 5,744 nt, median: 1006 nt). Gene Ontology enrichment analysis of putative *cis*-acting target genes in genomic proximity to lncRNAs revealed significant overrepresentation of functional categories essential for survival and dimorphic transition, including regulation of transcription by RNA polymerase II (5%), oxidoreductase activity (4.2%), and methylation (2%). In contrast, circRNAs demonstrated heterogeneous size distributions, with lengths ranging from 135 nt to 4,943 nt (median: 553 nt), predominantly clustering within the 200 bp to 1 kb range (77.1% of total circRNAs). Genomic distribution analysis revealed that 80.0% of identified circRNAs were derived from protein-coding gene loci, with derivation from genes involved in basic cellular processes such as translation (4%), intracellular protein transport (1%), cell redox homeostasis (0.6%). Although circRNAs demonstrated universally lower expression levels compared to protein-coding transcripts (Figure 1D), principal component analysis revealed that circRNA expression profiles effectively clustered samples according to morphological states and temporal progression (Figure 1E), indicating significant indicator or regulatory functions of circRNAs during dimorphic transition processes. This limited enrichment of *cis*-acting functions for circRNA host genes and lncRNA targets may suggest that these ncRNAs primarily exert their regulatory effects through trans-acting mechanisms rather than local genomic regulation.

### Dynamic transcriptional landscapes reveal stage-specific biological processes during *T. marneffei* dimorphic transition

Building upon the comprehensive landscape of transcriptomes, we next sought to delineate the specific biological processes occurring at distinct temporal phases during dimorphic transition. We employed the EBSeq-HMM to perform temporal module identification on protein-coding gene expression matrices across 15 time points for both mycelium-to-yeast and yeast-to-mycelium transitions^23^. Given the computational constraints of simultaneously analyzing 45 samples (memory requirements >800 GB), we strategically partitioned samples based on expression similarity patterns (Figure 1B). For M2Y transition, early time points (0, 0.5, 1, 2, 3 h) were designated as the M2Y_start group, with remaining time points (6-108 h) forming the M2Y_transition group, using 3 h as the comparative baseline. Similarly, for Y2M transition, time points 0–12 h comprised the Y2M_start group, while 24–108 h constituted the Y2M_transition group, with 12 h serving as the transition baseline. This temporal classification aligned remarkably well with morphological observations during cultivation and the notably slower initiation of Y2M compared to M2Y transition (Figure 1A and Figure S1), demonstrating tight correspondence between morphological and transcriptional dynamics.

Functional enrichment analysis of temporally regulated genes revealed distinct stage-specific biological programs. Consistent with sample clustering patterns, both M2Y (Figure 2A) and Y2M (Figure 3) transitions exhibited pronounced temporal staging in their enrichment profiles. During the M2Y_start phase, ATP-related energy pathways underwent extensive regulation, while sugar metabolism-associated energy pathways were activated during M2Y_transition, suggesting metabolic shifts corresponding to yeast morphology energy preferences. Concurrently, microtubule binding, DNA-related processes, and various biosynthetic pathways were activated during M2Y_start, whereas transcriptional and translational processes predominated during M2Y_transition. In contrast to M2Y, the Y2M process demonstrated greater complexity while maintaining similar temporal staging. Early Y2M phases involved extensive metabolic and biosynthetic changes, corresponding to highly active ribosomal biogenesis, transcription, translation, tRNA synthesis, and modification processes (Figure S3). During the Y2M_transition phase, when morphological examination revealed extensive asymmetric polarization from yeast cells generating hyphal structures, DNA-related pathways became activated, potentially reflecting the transition from uninucleate yeast cells to multinucleate hyphal forms.

**Figure 2.**
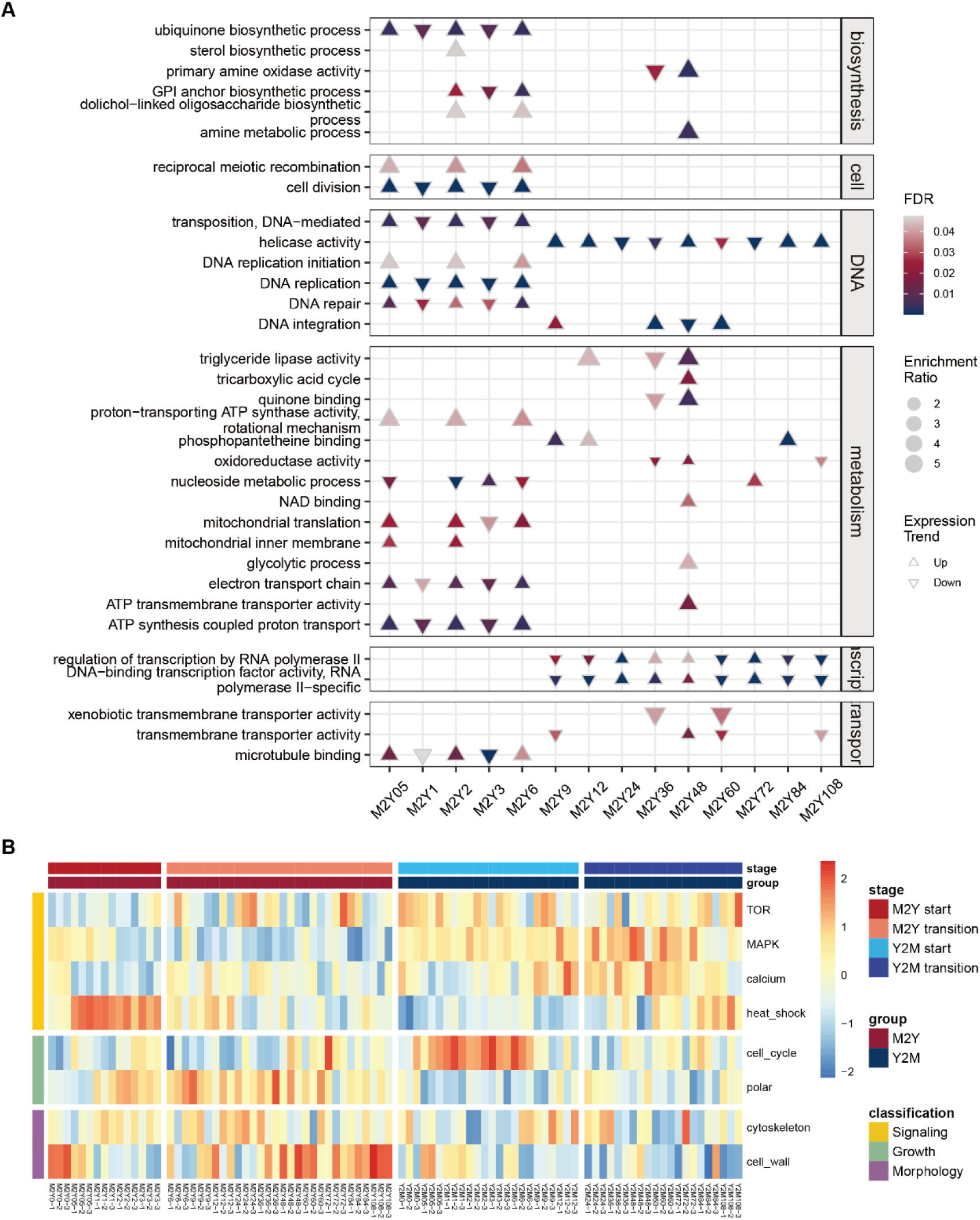
Functional profiling of transcriptomic reprogramming during M2Y transition in *T. marneffei*. (A) Gene Ontology (GO) enrichment analysis of differentially expressed genes (DEGs) at distinct time points of M2Y transition. Triangles denote gene expression trends, with upward triangles indicating upregulation and downward triangles revealing downregulation. Triangle size represents enrichment ratio, while color intensity reflects significance (FDR). Time points are annotated corresponding to transition phases. (B) Heatmap showing activation dynamics of signaling pathways, growth-related processes, and morphology-related functions during the dimorphic phase transition. Columns represent transcriptomic time points across M2Y and Y2M transitions, and colors indicate *Z*-score normalized expression values.

**Figure 3.**
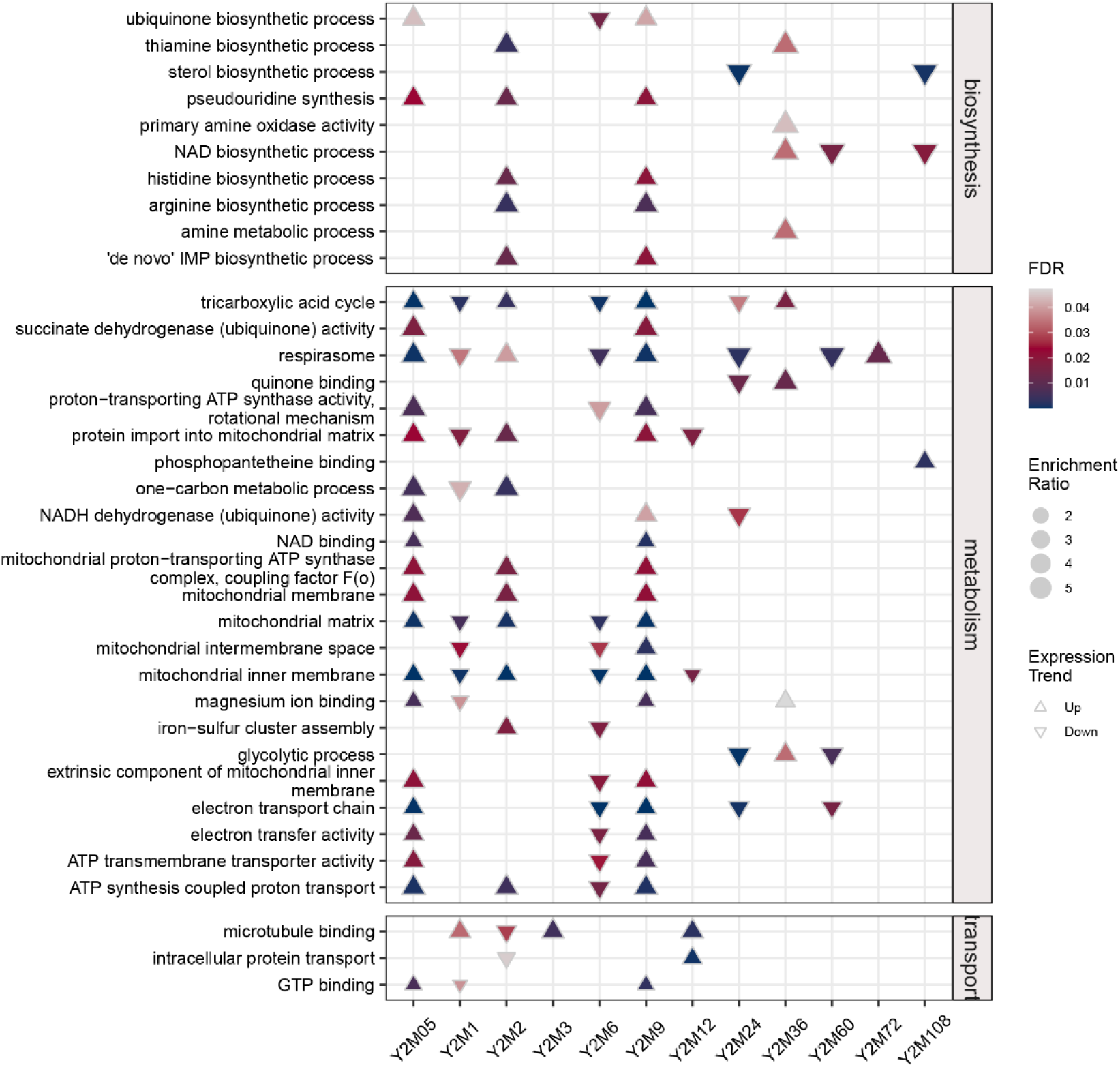
Functional insights into transcriptomic changes during Y2M transition in *T. marneffei*. GO-term enrichment analysis of DEGs during Y2M phase transition. Upward-pointing triangles represent upregulated genes, while downward-pointing triangles represent downregulated genes. Triangle size depicts enrichment ratio and color denotes FDR values. Time points are marked corresponding to transition phases.

**Figure S3.**
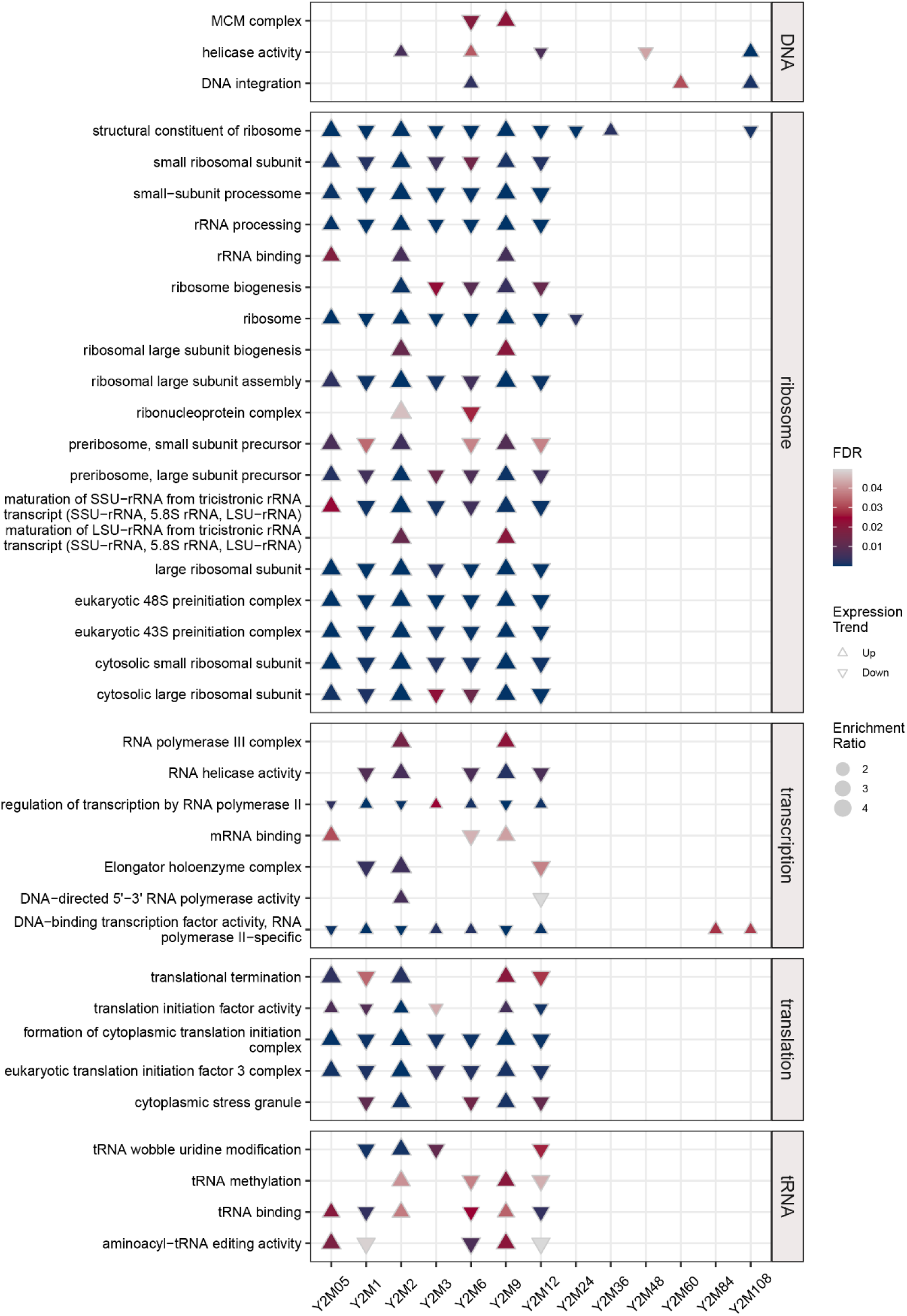
GO enrichment during Y2M transition in *T. marneffei*. GO-term enrichment analysis of DEGs during Y2M transition, visualized by expression trends, enrichment ratios, and FDR values over time points.

**Figure S4.**
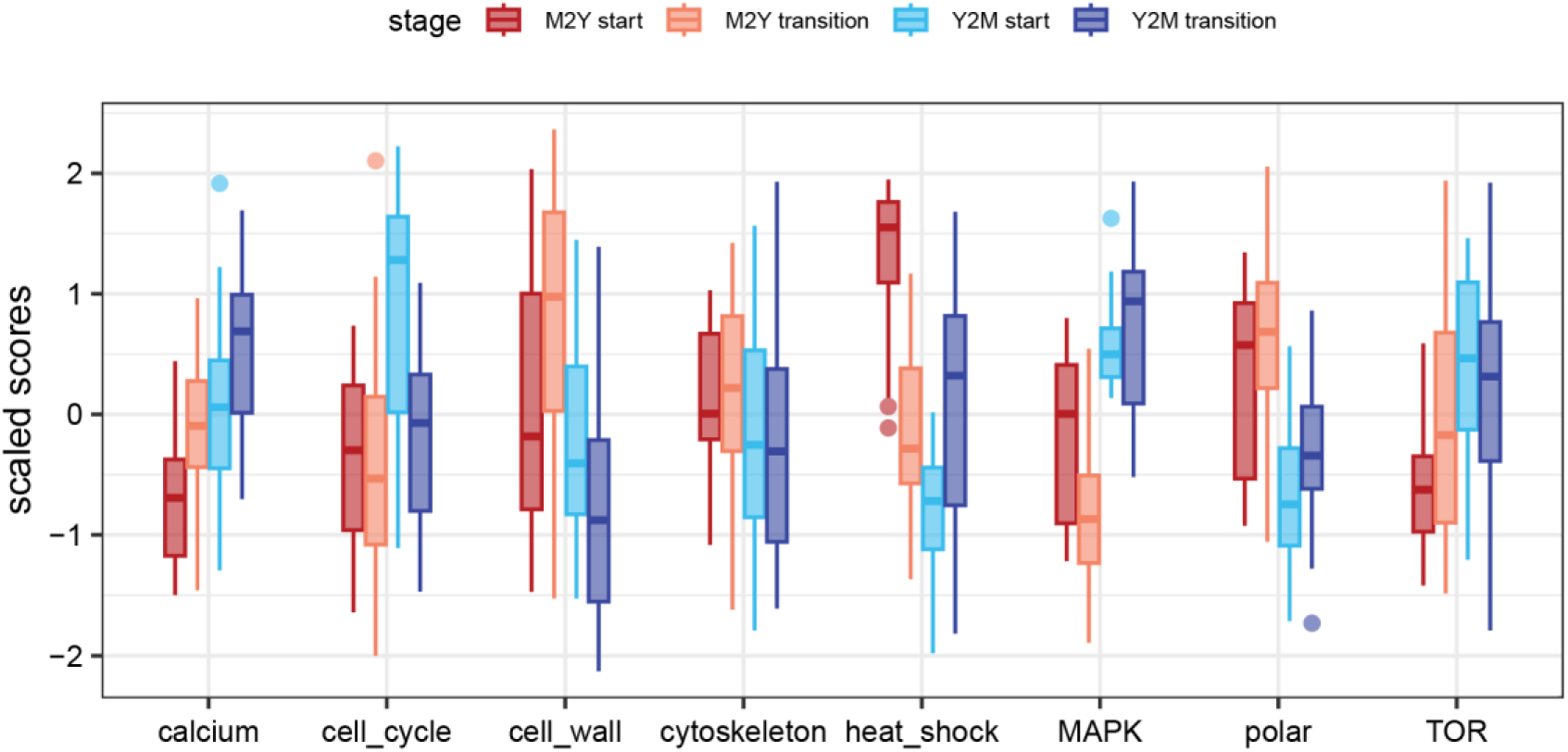
Stage-specific functional pathway activity during dimorphic transitions in *T. marneffei*. Boxplot visualization of scaled scores for key functional pathways across four distinct stages of dimorphic transition in T. marneffei. Each boxplot shows the distribution of scaled pathway activity scores, with the horizontal line inside the box representing the median and the box boundaries corresponding to the first quartile (Q1) and third quartile (Q3). The whiskers extend from the box to 1.5 times the interquartile range (IQR). Circles represent individual data points.

To capture fungal-specific morphological processes not reflected in standard GO enrichment, we defined eight custom gene sets based on functional relevance to dimorphic transition (Table S1). Normalized pathway GSVA scores revealed that M2Y transition exhibited prominent cell wall pathway activation during both earliest and latest phases, with intervening periods characterized by polar growth and cytoskeletal gene expression. Y2M transition displayed contrasting patterns, with early cell cycle gene activation followed by upregulation of multiple signaling pathways and polar growth, accompanied by significant downregulation of cell wall-related genes, consistent with compositional shifts between hyphal and yeast cell walls (Figure 2B & Figure S4).

Integration of detailed temporal transcriptional data enabled refined biological pathway mapping of *T. marneffei* dimorphic transition processes. During M2Y initiation, cell wall processes are first activated, followed by heat shock responses, substance transport, mitochondrial metabolism, and DNA-related processes, coinciding with hyphal fragmentation and arthroconidial formation. Subsequently, transcriptional processes, sugar metabolism, cytoskeletal dynamics, and cellular polarity respond coordinately, culminating in cell wall remodeling and yeast cell formation. Conversely, Y2M initiation involves activation of cell cycle-related genes alongside broad energy metabolism, substance transport, biosynthesis, transcription, and translation pathways, persisting for approximately 12 hours while maintaining yeast cell morphology until adequate material foundations for hyphal transition are established. This is followed by upregulation of multiple signaling pathways while maintaining transcriptional and metabolic activity levels until transformation into characteristic elongated hyphae.

### Cross-species heat-stress transcriptomics reveals a *T. marneffei*-specific thermoadaptive response program

Given that the preceding analysis revealed heat stress as a critical trigger for *T. marneffei* dimorphic transition, with specific temporal patterns of metabolic reprogramming and cellular remodeling, we next investigated whether these heat-responsive capabilities represent evolutionary adaptations unique to *T. marneffei* among *Talaromyces* species, by performing rRNA-depletion RNA-seq on closely related non-dimorphic species *Talaromyces stipitatus* and *Talaromyces amestolkiae* at 0, 1, and 3 hours post-heat stimulation. Using *Aspergillus fumigatus* as an outgroup, we identified 11,289 orthologous genes, including 4,082 single-copy orthologs. In contrast to the coordinated responses observed in *T. marneffei* at 1 and 3 hours, both *T. stipitatus* and *T. amestolkiae* exhibited substantial transcriptional divergence. Cosine distance analysis revealed that *T. amestolkiae* was more similar to *T. marneffei* than *T. stipitatus*, and the differences in gene expression were more pronounced than those caused by heat treatment (Figure 4A), indicating the pre-existing differences in transcriptional background among the three species.

**Figure 4.**
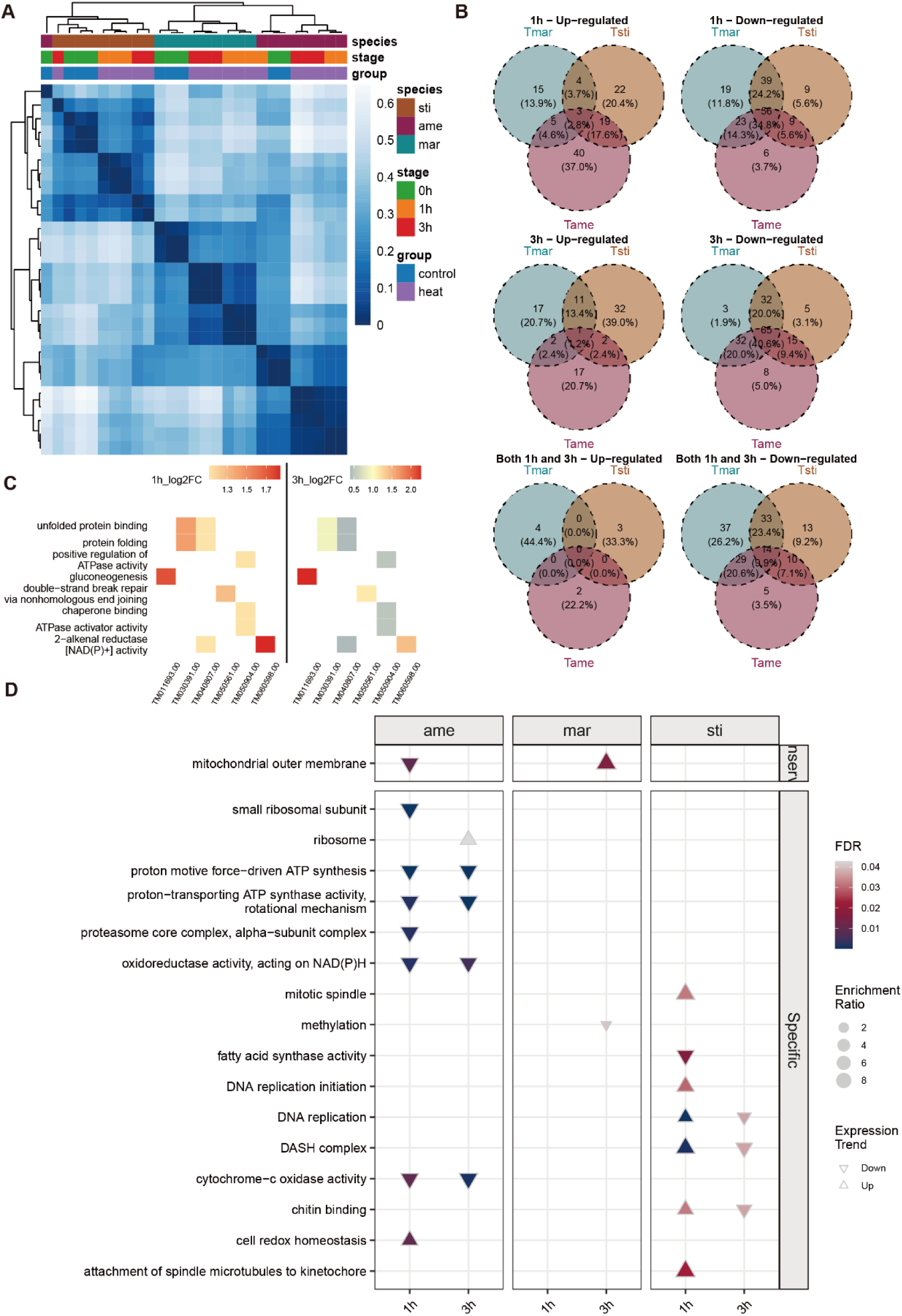
Comparative transcriptomic responses to heat stress across *Talaromyces* species. (A) Heatmap of transcriptomic patterns across three *Talaromyces* species under control and heat stress conditions at 0 h, 1 h, and 3 h. Rows and columns represent species and conditions hierarchically clustered based on Euclidean distances. Color intensity signifies log2 transformed fold-change values, with darker blue for lower expression and lighter blue for higher. (B) Venn diagrams illustrating overlaps of upregulated (left) and downregulated (right) genes among species at 1 h and 3 h post-heat stress. Numbers indicate gene counts, and percentages in parentheses indicate relative proportions. (C) Heatmap visualizing enriched GO terms for DEGs uniquely expressed orthologs at 1 h and 3 h in *T. marneffei*. Dot color represents log2(FoldChange) values, while dot size denotes enrichment significance (FDR). (D) Analysis of species-specific GO enrichment under heat stress at 1 h and 3 h. Upward triangles represent upregulated genes, while downward triangles signify downregulated genes. Triangle size and color scale correlate with enrichment ratio and FDR values.

Comparative ortholog expression analysis also revealed species-specific heat response strategies in *Talaromyces*. Although similar to the heat response pattern of *T. marneffei*, *T. amestolkiae* and *T. stipitatus* exhibited substantial proportions of differentially upregulated orthologs at 1 and 3 hours, no orthologs yet were commonly upregulated across all three species (Figure 4B). This suggests that while *T. amestolkiae* and *T. stipitatus* possess copies of genes upregulated in *T. marneffei*, their protective gene expression programs in response to heat stimulation differ fundamentally. Notably, we identified 15 orthologous genes consistently upregulated in *T. marneffei* at both 1 and 3 hours while being downregulated in *T. amestolkiae* and *T. stipitatus*, including transcription factor gene *TM021177.00*. These inversely regulated genes exhibited significant enrichment in metabolic processes and protein folding pathways (Figure 4C), corroborating the temporal expression patterns observed at M2Y initiation. Such convergent evidence indicates that targeted upregulation of these pathways likely constitutes a prerequisite for developing thermoadaptive capacity.

To directly examine whether heat-responsive genes overlap with dimorphic transition regulators, we compared the 15 *T. marneffei*-specific upregulated orthologs with genes that exhibited differential expression at more than three time points during the M2Y transition. Notably, 14 of the 15 genes demonstrated significant overlap, indicating that the heat stress response machinery has been co-opted for dimorphic transition regulation. In contrast to the upregulation pattern, over 30% of downregulated genes exhibited consistent expression trends among differentially expressed orthologs at both time points, with the highest proportion of shared homologs observed at 3 hours. Notably, no orthologs were identified that showed downregulation in *T. marneffei* concurrent with upregulation in other species, further supporting the evolutionarily distinct positive regulatory circuitry that underpins the thermoadaptive transcriptional program of *T. marneffei*.

We also wonder whether the same transcriptional changes could be found in the pathway levels. Pathway-level analysis corroborated the transcriptional divergence observed in ortholog expression patterns (Figure 4D). While *T. marneffei* significantly downregulated methylation processes and upregulated cellular respiration, *T. amestolkiae* strongly suppressed cellular respiration, potentially reducing mitochondrial activity to prevent reactive oxygen species burst, though this strategy may compromise growth efficiency. *T. stipitatus* upregulated cell division-related pathways, suggesting attempts to accelerate proliferation for rapid environmental adaptation, but potentially increasing cellular damage risk due to insufficient protective mechanism activation. Importantly, no pathways exhibited shared differential trends among all three species, consistent with ortholog analysis results and further reflecting distinct survival strategies under heat stress.

### Comparative genomics of divergent response genes uncovers *cis*-regulatory modules underlying species-specific heat responses

To elucidate the mechanistic basis of transcriptomic differentiation, we systematically analyzed promoter sequence characteristics underlying differential gene expression patterns. Through *de novo* motif discovery and comparative analysis of directionally regulated gene sets, we identified conserved and species-specific transcriptional regulatory signatures. Motif enrichment analysis revealed five primary transcription factor classes governing differential expression: C2H2 zinc finger factors, C6 zinc cluster factors, heteromeric CCAAT-binding factors, RING-type zinc fingers, and TATA-binding proteins (Figure 5A). A conserved regulatory module emerged across all three species, characterized by significant enrichment of AZF1 binding motifs in downregulated gene promoters (*p* = 1.71×10⁻⁵ to 6.99×10⁻⁵), indicating universal involvement of this C2H2 zinc finger factor in stress-induced transcriptional repression.

**Figure 5.**
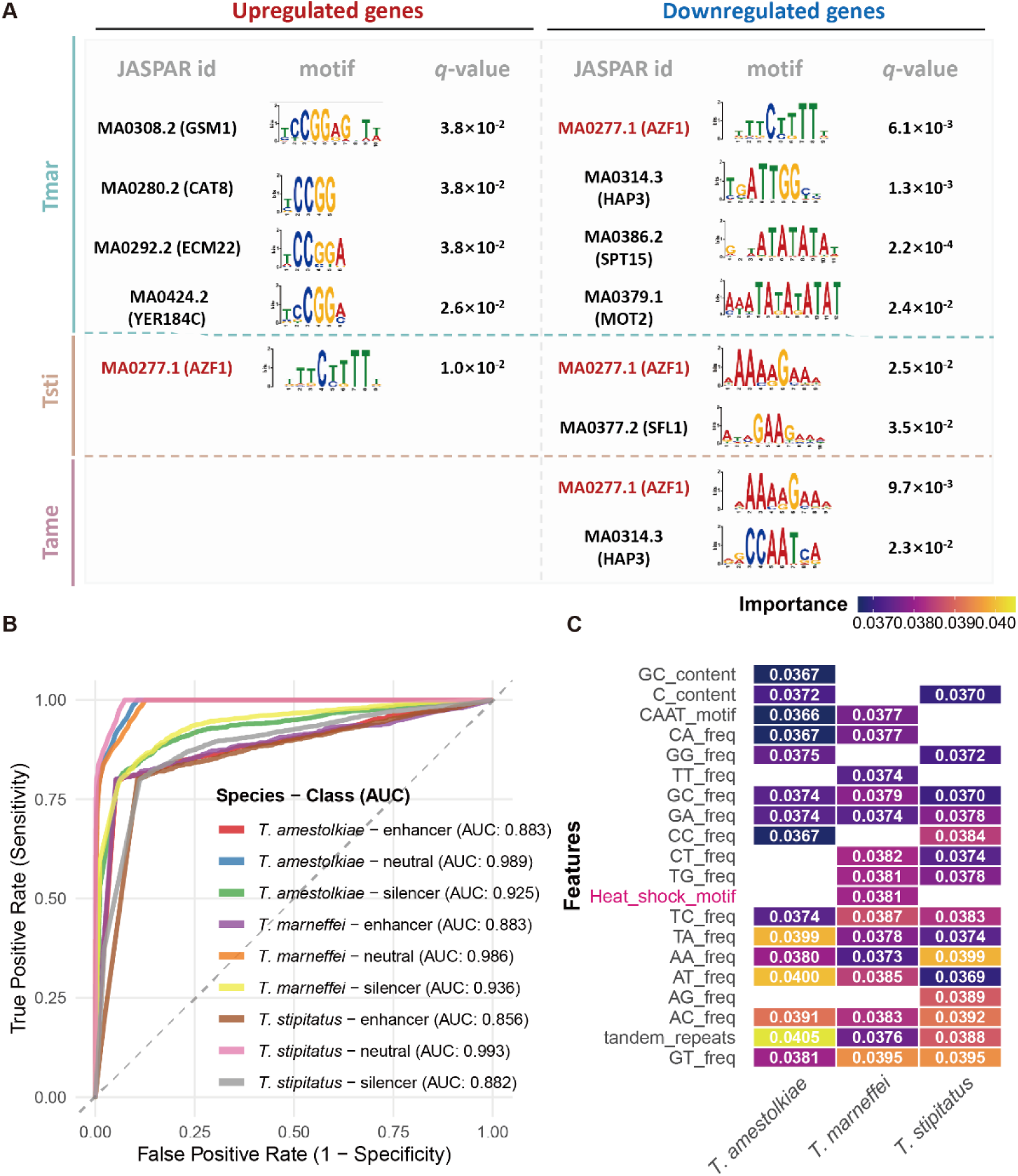
Regulatory signatures and predictive modeling of promoter regions in *Talaromyces* species. (A) Transcription factor motif enrichment analysis corresponding to upregulated (left) and downregulated (right) DEGs across *T. marneffei*, *T. stipitatus*, and *T. amestolkiae*. Each panel showcases identified motifs with JASPAR IDs, representative logos, and statistical significance (*q*-values). (B) Receiver operating characteristic (ROC) curves demonstrating the performance of predictive models for identifying enhancer, silencer, and neutral regions in the three species, with area under curve (AUC) metrics. The dashed diagonal line illustrates the performance of a random classifier. (C) Feature importance analysis for machine learning classifiers in (B), displayed as heatmaps. Color intensity indicates feature contributions to predictive performance across species.

Species-specific regulatory patterns provided insights into evolutionary divergence in stress response mechanisms. The heteromeric CCAAT-binding factor HAP3 motif (consensus: TGATTGGCT) was preferentially enriched in *T. amestolkiae* and *T. marneffei* downregulated genes (*p* = 1.32×10⁻⁴ and 3.68×10⁻⁶, respectively), suggesting conserved roles in cell cycle and metabolic regulation, which is consistent with the pathway enrichment. Conversely, *T. stipitatus* exhibited unique enrichment of heat shock factor SFL1 motifs (consensus: ATAGAAGAAA, *p* = 1.98×10⁻⁴) in downregulated genes, indicating alternative stress response pathway activation. Notably, *T. marneffei* exhibited distinctive upregulation patterns characterized by coordinated enrichment of C6 zinc cluster transcription factor binding motifs (CAT8, ECM22, GSM1, SIP4, YER184C; *p*-values ranging from 5.24×10⁻⁴ to 7.67×10⁻⁴), which predominantly recognize TCCGG/TCCGGA motif variants. This enrichment of C6 zinc cluster transcription factor regulatory elements corresponding to their transcriptional activation, providing a mechanistic basis for the observed upregulation of multiple energy and metabolic pathways. Furthermore, TATA-binding protein SPT15 motifs (consensus sequence: ATATATATATC, p = 6.23×10⁻⁶) and RING-type zinc finger MOT2 motifs (p = 2.78×10⁻⁴) were exclusively detected in *T. marneffei* downregulated genes, indicating coordinated repression of core transcriptional machinery and protein degradation pathways.

To comprehensively characterize promoter regulatory potential beyond binding motifs, we employed Random Forest classification using sequence composition features including dinucleotide/trinucleotide frequencies, GC content, repetitive elements, and canonical regulatory motifs. Models achieved robust classification performance (86.7-91.7% accuracy) with low standard deviation (SD < 0.01) in 5-fold cross-validation, indicating stable predictive capacity across regulatory categories, including enhancers (from up-regulated genes under heat), silencers (from down-regulated genes under heat), and neutral elements (from non-significantly differential expressed genes under heat) (Figure 5B). Feature importance analysis revealed profound species-specific regulatory architectures (Figure 5C). *T. marneffei* exhibited a GT dinucleotide-dominated signature (importance: 0.0395), with heat shock motifs uniquely ranking among top predictive features (0.0381), reinforcing its specialized thermal stress adaptation. *T. stipitatus* displayed fundamentally distinct purine-rich regulation through AA dinucleotide prominence (0.0399) and significant tandem repeat contributions (0.0388), suggesting repetitive element-mediated regulatory mechanisms. *T. amestolkiae* demonstrated tandem repeat-dominated architecture (0.0405) coupled with AT-rich signatures (AT: 0.0400; TA: 0.0399), potentially conferring distinct chromatin organization and heat sensitivity profiles. These findings collectively indicate that transcriptomic divergence among *Talaromyces* species is associated with differences in promoter architecture and transcriptional regulatory networks, which may partially explain the evolutionary basis of thermotolerance of *T. marneffei*.

### *Trans*-regulatory lncRNAs and circRNAs complete the regulatory repertoire of dimorphic-transition pathways

To uncover how *trans*-acting non-coding RNAs orchestrate the transcriptomic rewiring underlying thermal dimorphism, we employed Graphical Gaussian Models (GGMs) by integrating circRNAs, lncRNAs, and protein-coding genes across temporal dimorphic transitions to elucidate potential trans-acting mechanisms. Network topology analysis revealed significant structural reorganization between the initiation and transition phases, where lncRNA hubs predominated during early response phases such as M2Y_start and Y2M_start, whereas circRNA hubs became more prominent during the morphological transition phases (Figure S5). This temporal stratification suggests distinct regulatory mechanisms whereby lncRNAs establish initial transcriptional programs while circRNAs fine-tune transition-specific processes.

During M2Y initiation (Figure 6A), network analysis identified a significant hierarchical regulatory cascade coordinating cell wall remodeling and metabolic reprogramming, aligning with the observation that these are the primary pathways altered during the M2Y transition. Core melanin biosynthesis genes (*TM011794.00*: polyketide synthase, *TM060719.00*: scytalone dehydratase, *TM020808.00*: cutinase) formed tightly interconnected modules directly regulated by circRNA pm1_01:1603420|1604411, which is derived from F-box protein *TM010590.00*. Concurrently, aspergillopepsin F precursor (*TM051145.00*) anchored an extensive regulatory network that sequentially coordinated signal perception through chemokine receptor-like 2, followed by signal transduction via G-protein regulatory factor egl-10 and associated lncRNAs, which then regulated metabolic processes including protein degradation and carbon uptake, subsequently controlling transport systems for sugar and pantothenate, and finally governing membrane remodeling and secondary metabolism.

**Figure 6.**
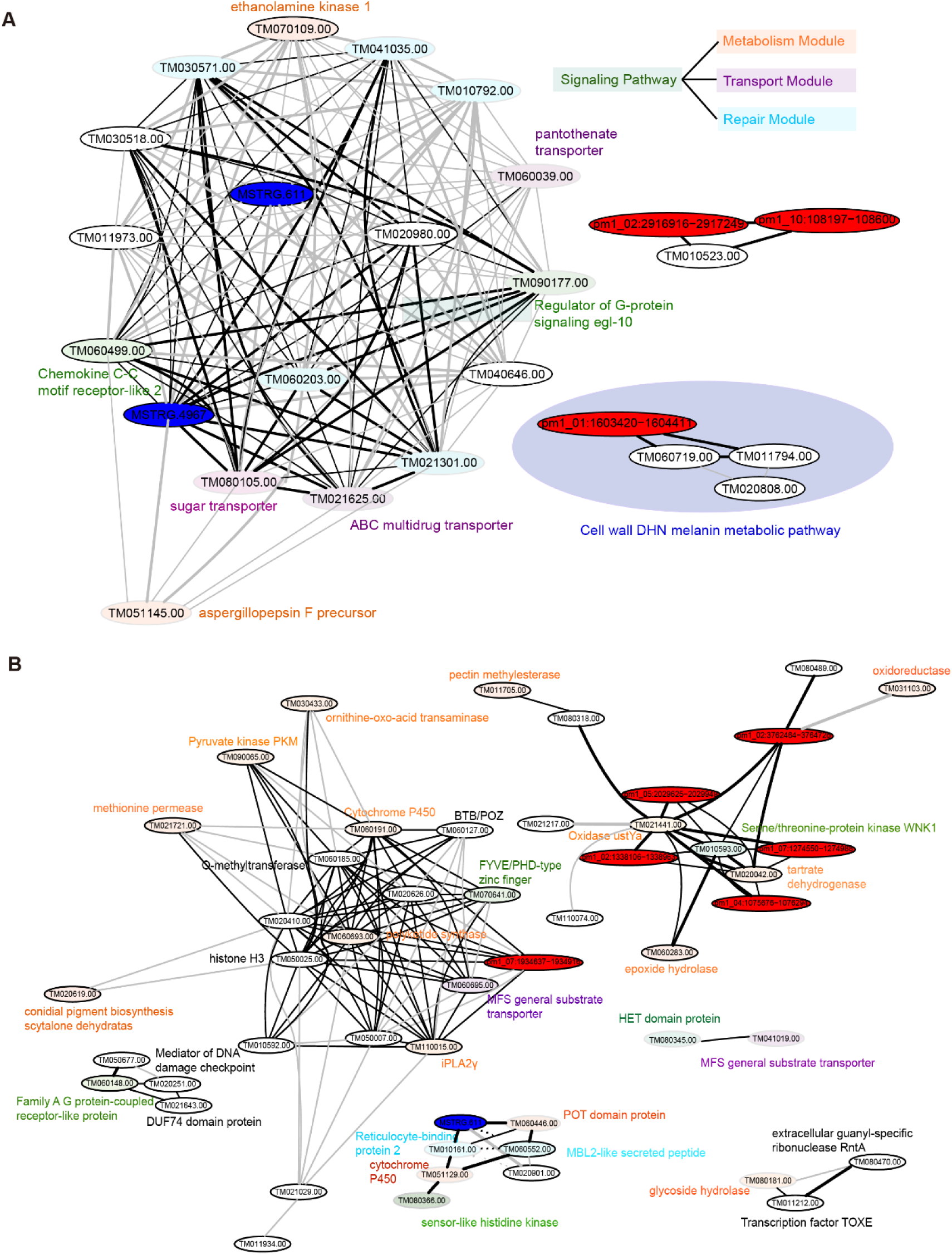
Co-expression networks of ncRNAs underlying dimorphic transitions in *T. marneffei*. (A) Visualization of key gene co-expression networks during the M2Y transition. Nodes represent individual genes, with edges denoting significant co-expression relationships. Edge thickness reflects correlation strength, and nodes are color-coded by predicted functional modules: red, metabolism; purple, transport; blue, repair; green, signaling pathways. (B) Co-expression network of key genes during the Y2M transition, with functional roles annotated by node color.

**Figure S5.**
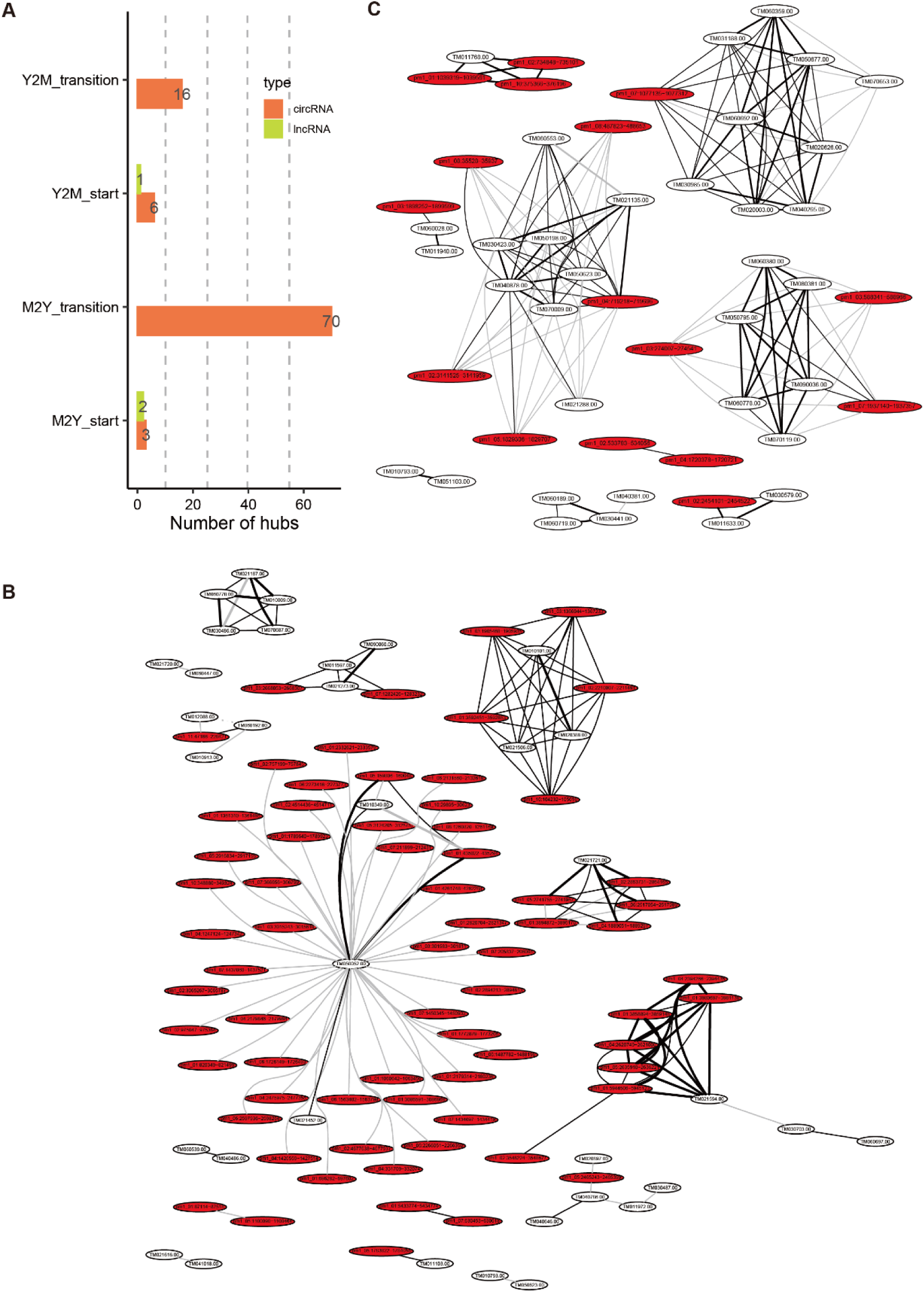
CircRNA hubness and network associations during dimorphic transitions. (A) Bar plot showing the number of hub nodes for circRNA (orange) and lncRNA (green) identified in four experimental conditions: Y2M_start, Y2M_transition, M2Y_start, and M2Y_transition. (B-C) Co-expression networks of circRNAs and their gene associations during M2Y_transition (B) and Y2M_transition (C), with hub circRNAs highlighted (red nodes) and edge thickness indicating co-expression strength.

Y2M initiation networks (Figure 6B) demonstrated distinct organizational principles, with lncRNA MSTRG.611 serving as a central coordinator linking environmental sensing (*TM080366.00*: histidine kinase), transcriptional control (*steA*), metabolic redistribution (Cytochrome P450, POZ proteins), and structural remodeling (*Rbp2*, MBL2-like peptides). Notably, early-response circRNA pm1_07:1934637|1934916 exhibited regulatory associations with histone H3-mediated chromatin modifications, potentially coordinating epigenetic control of key metabolic genes through acetylation and demethylation mechanisms. Serine/threonine kinase WNK1 (*TM010593.00*) emerged as an additional metabolic hub directly associated with multiple circRNAs, reinforcing the central role of ncRNA-mediated kinase regulation in morphological transitions.

### Integrated genomic-transcriptomic comparison reveals distinct evolutionary positions of ncRNAs in dimorphism and heat response

The observation that ncRNAs coordinate complex species-specific transcriptional programs necessitated investigation of their evolutionary origins and conservation patterns. To determine whether identified these ncRNAs reflect species-specific adaptations or conserved regulatory mechanisms, we conducted comprehensive comparative genomic and transcriptomic analyses across nine *Talaromyces* species.

Genomic sequence alignment revealed striking disparities between conservation at genomic versus transcriptomic levels. Among lncRNAs, 63 (26.3%) showed genomic conservation across multiple species, including four conserved across five species (Figure S6). However, transcriptomic conservation proved markedly restricted with only four lncRNAs demonstrated expression conservation, with none exhibiting pan-species expression patterns. This genomic-transcriptomic discordance was less pronounced for circRNAs. A total of 2,568 circRNAs (2,519 gene-derived, 49 intergenic) were identified as genomically conserved, in which 59 maintained transcriptomic conservation (Figure 7A). Species-specific distribution patterns revealed hierarchical conservation, with 39 circRNAs homologous to *T. amestolkiae*, 15 to *T. stipitatus*, and merely 5 shared across all three species (Figure 7B). Moreover, the five transcriptomically conserved circRNAs exhibited significantly elevated expression specifically in *T. marneffei* (Figure 7C) with pronounced temporal regulation during dimorphic transition (Figure 7D). Network analysis uncovered extensive regulatory divergence despite sequence conservation. Conserved circRNAs (consCirc2, consCirc4) predominantly interacted with *T. marneffei*-specific expressed orthologous genes, with only consCirc3 retained pan-species ortholog interactions (Figure 7E). These findings indicate that despite sequence conservation, some circRNAs have undergone functional rewiring in *T. marneffei* as the as protein-coding genes, demonstrating evolutionary co-option of conserved sequences for species-specific regulatory innovation.

**Figure 7.**
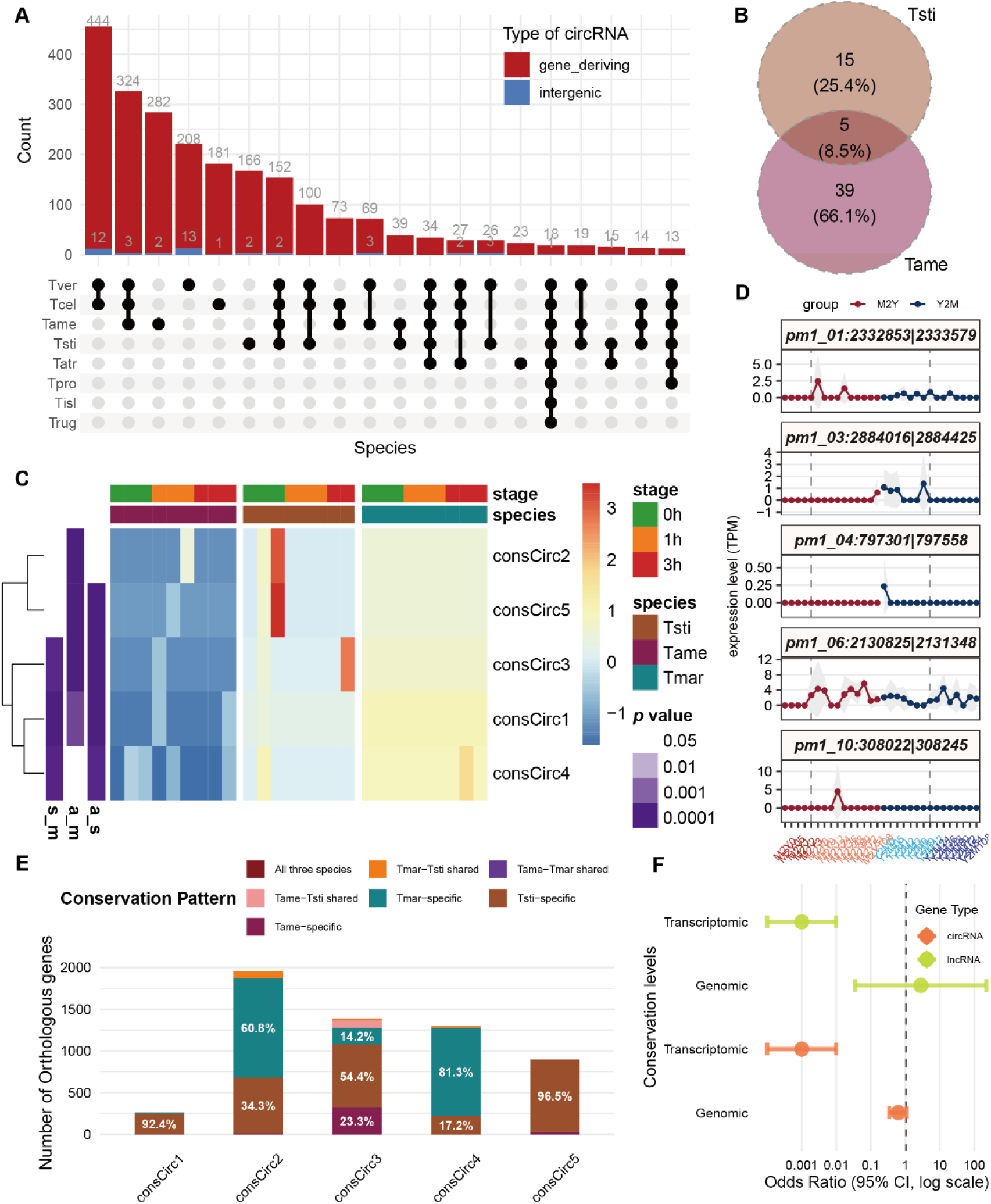
Conservation and functional dynamics of ncRNAs in *Talaromyces* species. (A) Genomic conservation of circRNAs across nine *Talaromyces* species, with classification into gene-derived (red) and intergenic (blue) categories. The bar chart above shows circRNA converd with *T. marneffei* counts per species, while the bottom matrix reveals species-specific overlaps in circRNA presence. (B) Venn diagram illustrating the overlap of circRNAs conserved with *T. marneffei* identified in the transcriptomes of *T. stipitatus* and *T. amestolkiae*. (C) Heatmap of normalized expression levels of five conserved circRNAs across post-heat stages (0 h, 1 h, 3 h) and species. *Z*-scores represent standardized expression. (D) Line plots depicting time-course expression (TPM) dynamics for five conserved circRNAs in *T. marneffei*. The x-axis represents time points, while the y-axis indicates mean TPM values across biological replicates (mean ± SD shown in gray shading). (E) Comparison of orthologous target gene numbers trans-interacting with the five conserved circRNAs. (F) Forest plot of conservation odds for non-coding RNAs serving as network hubs.

**Figure S6.**
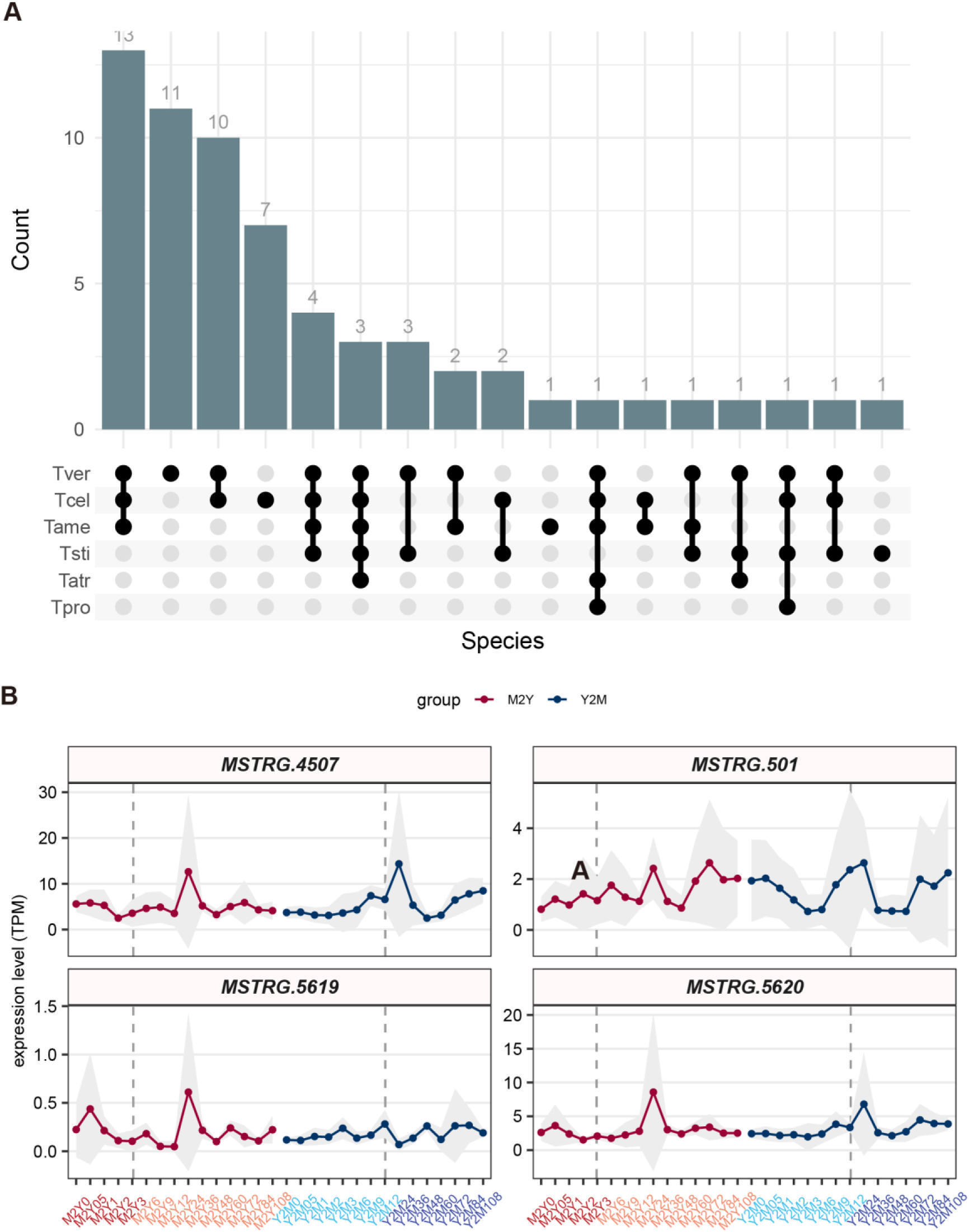
Conservation and expression dynamics of lncRNAs in *Talaromyces* species. (A) Genomic-conserved lncRNA identification across nine Talaromyces species. The bar chart above highlights the total count of conserved lncRNAs per species, with the bottom matrix showcasing the overlap of circRNA presence across species. (B) Line plots detailing TPM-based time-series expression trends for four conserved lncRNAs in *T. marneffei*.

Systemic analysis further revealed contrasting evolutionary strategies between ncRNA classes. Genomically conserved lncRNAs demonstrated positive correlation with network hub roles (Fisher’s exact test, OR > 1, *p*-value < 0.05), indicating that sequence-conserved lncRNAs preferentially occupy central regulatory positions. Conversely, genomically conserved circRNAs exhibited negative correlation with hub status (Fisher’s exact test, OR < 1, *p*-value < 0.05). This dichotomy suggests that regulatory innovation operates through distinct mechanisms: lncRNAs maintain conserved architectural roles across species, whereas circRNAs achieve functional diversification through species-specific emergence rather than ancestral sequence modification.

To elucidate the mechanisms underlying species-specific circRNA generation, we investigated whether their biogenesis correlates with variations in RNA-binding protein (RBP) motifs within flanking sequences^24^. Through *de novo* motif discovery using STREME with exonic background correction, followed by cross-referencing with RBP databases (CISBP-RNA, RNAcompete, and YEAST), we identified significant enrichment of distinctive RBP motifs unique to circRNAs in 28.9% of *T. marneffei*, 54.0% of *T. amestolkiae* and 4.8% of *T. stipitatus* candidates, with stringent *q*-values below 0.05. Each species showed distinct enrichment patterns. In *T. marneffei*, the most enriched motif was RNAcompete|RNCMPT00059:REF2, whose corresponding protein likely participates in stress granule formation through RNA binding. *T. amestolkiae* showed predominant enrichment of RNAcompete|RNCMPT00078:TRA2, a motif associated with pre-mRNA splicing regulation, while *T. stipitatus* exhibited highest enrichment for CISBP|M148_0.6:PABPN1, corresponding to a nuclear poly(A)-binding protein. These distinct enrichment patterns indicate that different RBPs may govern circRNA biogenesis across these species. Further supporting the role of RBP motifs in circRNA evolution, we found that transcriptionally conserved circRNAs in *T. marneffei* showed markedly higher propensity for harboring identical RBP motifs compared to their non-conserved counterparts. Specifically, 33.8% of transcriptionally conserved circRNAs contained fully concordant RBP motifs, a significant enrichment relative to non-conserved circRNAs (Fisher’s Exact Test, two-sided, *p*-value = 2.95 × 10⁻³⁸). Among the shared motifs in conserved circRNAs, RNAcompete|RNCMPT00078:TRA2 was most prevalent, appearing in 65% (13/20) of cases, suggesting preserved RBP-mediated mechanisms under functional constraints. These findings align with previously proposed hypotheses suggesting that species-specific circRNA expression arises from accumulated genomic variations that alter splicing regulation. This evolutionary divergence may confer functional specialization, enabling circRNAs to acquire species-specific regulatory roles through differential RBP-mediated processing.

### Functional validation reveals evolutionary incorporation of ncRNAs into regulatory-network plasticity

To validate the functional roles of the aforementioned ncRNAs in *T. marneffei* morphic transition, we performed overexpression experiments on two *T. marneffei*-specific hub lncRNAs (MSTRG.611 and MSTRG.4967) and one *T. marneffei*-specific hub circRNA (M2Y_start: pm1_01:1603420|1604411) to investigate their regulatory roles in morphic regulation. Because direct quantification of the morphological transition relies primarily on subjective microscopic observation, we instead evaluated stress sensitivity assays as quantitative physiological proxies for morphological adaptability.

Ectopic overexpression yielded five MSTRG.611 mutant strains with significantly elevated transcript levels (Figure 8A). Although MSTRG.611 overexpression did not alter vegetative growth on SDA medium (Figure S7A), these mutants exhibited enhanced resistance to multiple stressors. At 25°C, the strains showed increased tolerance to SDS and NaNO_2_ (Figures 8B–C). A consistent stress-resistant phenotype was observed at 37°C, including heightened tolerance to SDS, Congo Red, and Menadione albeit at lower threshold concentrations (Figure 8D and Figures S7B–D). These findings align with the high centrality of MSTRG.611 in the Y2M network, particularly its strong regulatory associations with cell wall remodeling modules. To evaluate functional conservation, we also overexpressed MSTRG.611 in *T. stipitatus*. Despite the conservation of several interacting orthologs between *T. marneffei* and *T. stipitatus*, this overexpression yielded entirely opposite phenotypic effects, with the *T. stipitatus* mutants exhibiting a significantly reduced colony size on Congo Red (Figures S7E). This species-specific divergence highlights that while core regulatory modules may be conserved, the functional output of non-coding RNAs can rapidly adapt to serve lineage-specific physiological requirements.

**Figure 8.**
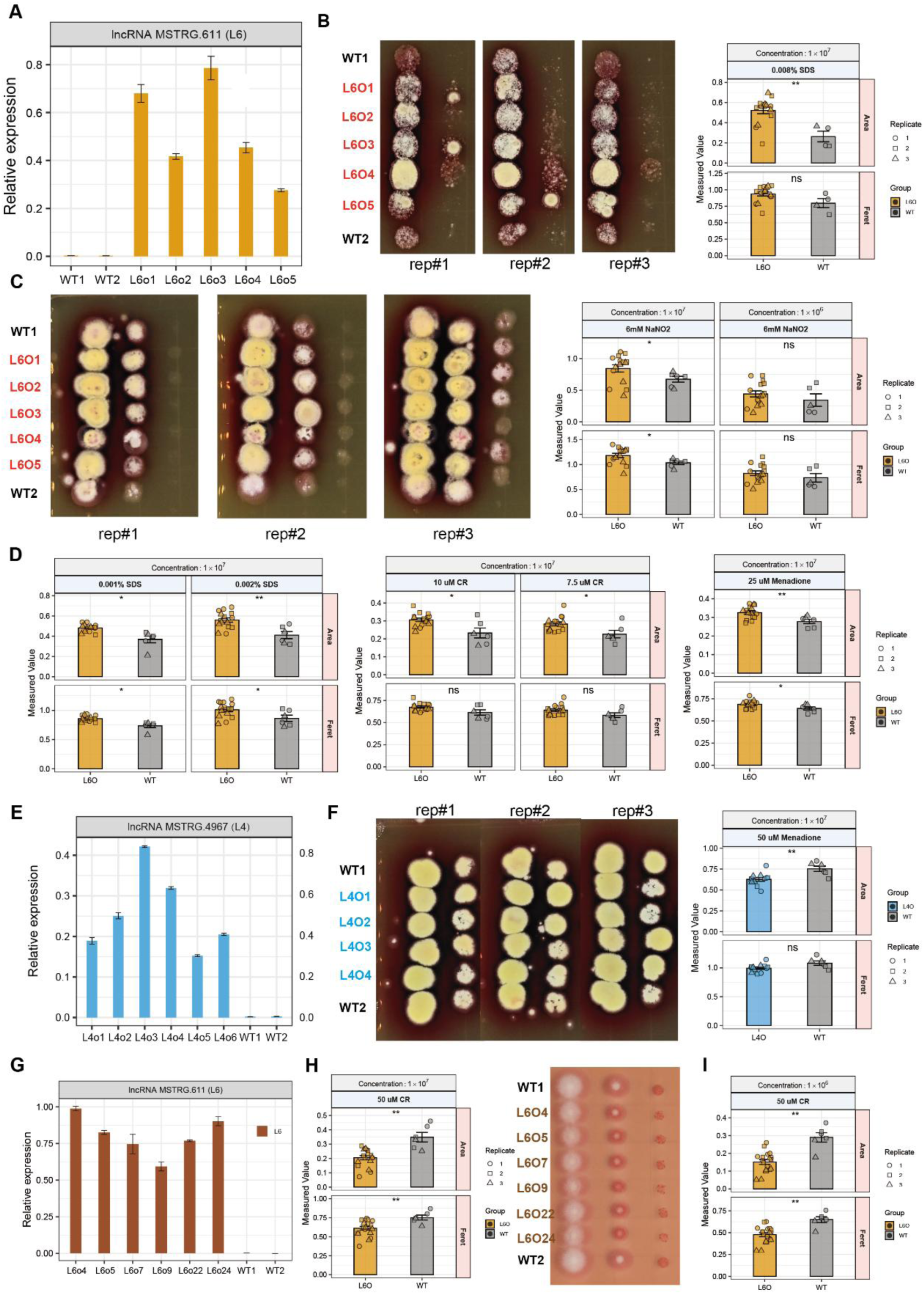
Functional characterization of species-specific lncRNAs MSTRG.611 and MSTRG.4967 in *T. marneffei*. (A) Quantitative RT-PCR (qRT-PCR) analysis of MSTRG.611 expression levels in two wild-type (WT) strains of *T. marneffei* and five independent ectopic overexpression (OE) mutant strains (L6O1–L6O5). Data represent mean ± SD. (B–C) Growth and stress tolerance assays at 25°C. Representative spot assays (left) and quantitative analysis of colony area and Feret diameter (right) showing the response of WT and L6O strains to (B) 0.008% SDS and (C) 6 mM NaNO_2_. (D) Quantitative analysis of stress resistance at 37°C. Bar plots illustrate the significantly enhanced tolerance of MSTRG.611 overexpression strains to cell wall stressors, including 0.001%–0.002% SDS and 7.5–10 μM Congo Red (CR), as well as oxidative stress induced by 25 μM Menadione. (E) Relative expression levels of lncRNA MSTRG.4967 in WT of *T. marneffei* and six OE mutant strains (L4O1–L4O6). (F) Growth and sensitivity assays of MSTRG.4967 OE strains at 25°C under 50 μM Menadione treatment. Representative spot assays (left) and quantitative analysis (right) indicate a significant increase in sensitivity to oxidative stress. (G) Quantitative RT-PCR (qRT-PCR) analysis of MSTRG.611 expression levels in two WT strains of *T. stipitatus* and six independent ectopic overexpression (OE) mutant strains. Data represent mean ± SD. (H) Quantitative analysis (left) and representative spot assays (right) of stress resistance at 25°C. Bar plots illustrate the significantly enhanced tolerance of MSTRG.611 overexpression strains of *T. stipitatus* to 50 μM Congo Red (CR). (I) Quantitative analysis of stress resistance at 37°C. Bar plots illustrate the significantly enhanced tolerance of MSTRG.611 overexpression strains of *T. stipitatus* to 50 μM Congo Red (CR). Statistical significance was determined using Student’s t-test (* *p* < 0.05, ** *p* < 0.01; ns, not significant). For all quantitative plots, individual data points from three biological replicates are displayed.

**Figure S7.**
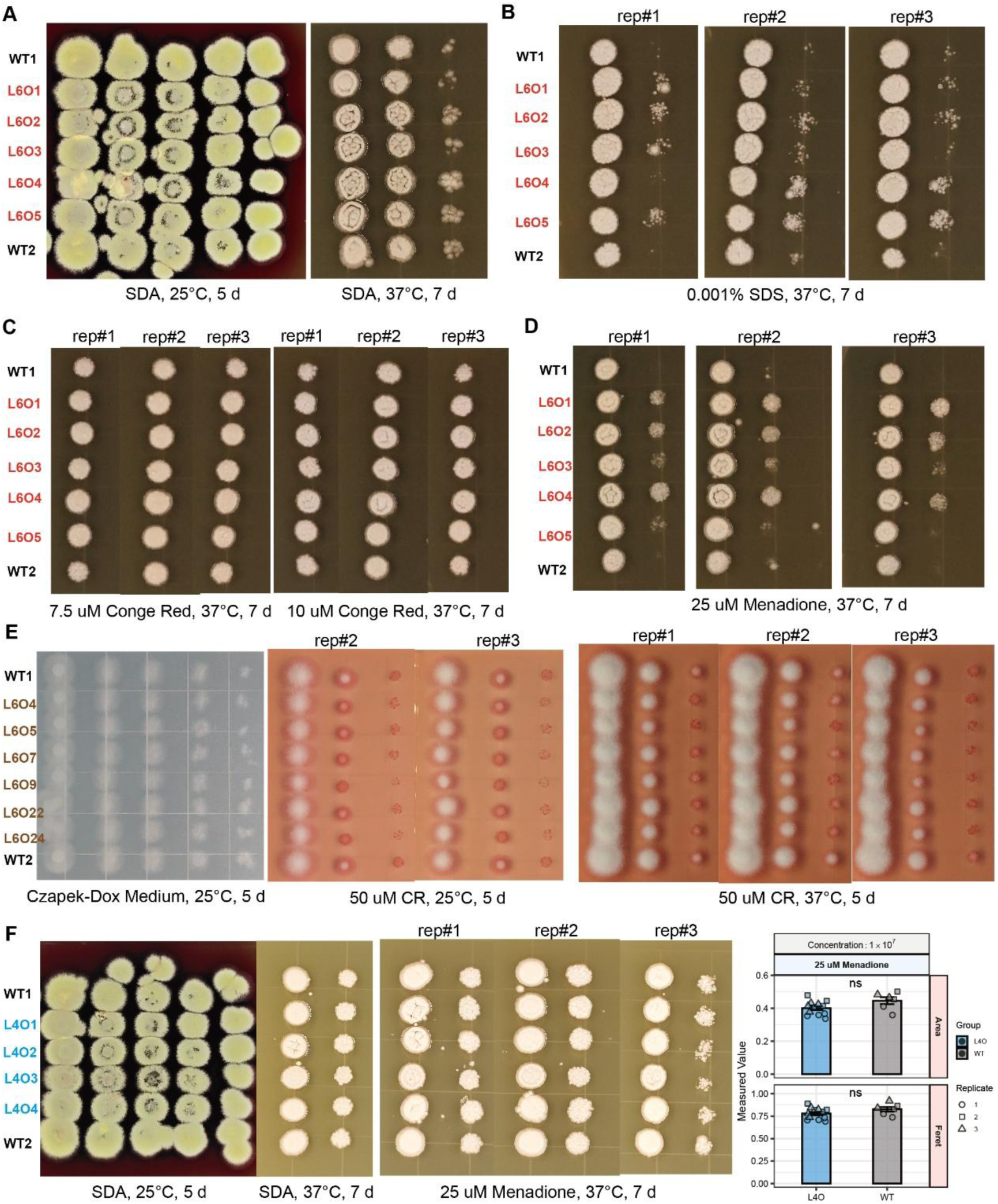
Phenotypic profiling of lncRNA overexpression strains under vegetative and stressful conditions. (A) Growth phenotypes of MSTRG.611 OE strains on standard SDA medium at 25°C (5 d) and 37°C (7 d), showing no significant difference in vegetative growth compared to WT. (B–D) Representative spot assays of MSTRG.611 OE strains at 37°C under treatment with (B) 0.001% SDS, (C) 7.5–10 μM Congo Red (CR), and (D) 25 μM Menadione. (E) Phenotypic analysis of MSTRG.611 OE strains of *T. stipitatus*. Left panels show normal vegetative growth on Czapek-Dox medium at 25°C and 37°C. Right panels show representative spot assays and quantitative analysis under 37°C. Quantitative data represent the mean of three biological replicates. Error bars indicate SD. (F) Phenotypic analysis of MSTRG.4967 OE strains. Left panels show normal vegetative growth on SDA at 25°C and 37°C. Right panels show representative spot assays and quantitative analysis under 25 μM Menadione at 37°C. Quantitative data represent the mean of three biological replicates. Error bars indicate SD.

**Figure S8.**
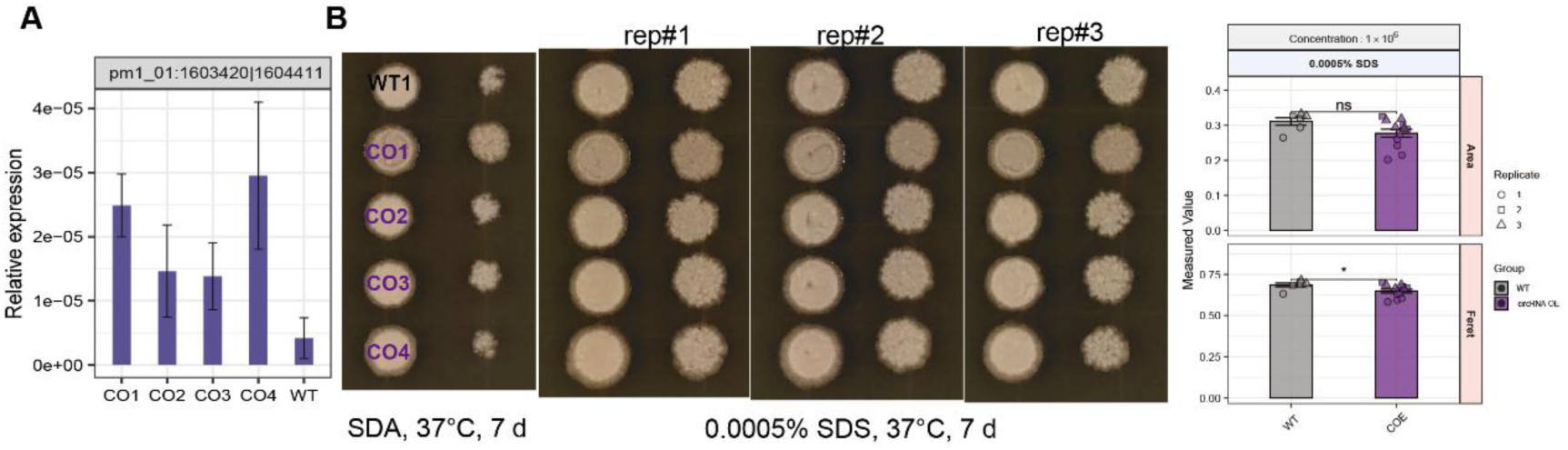
Phenotypic profiling of circRNA pm1_01:1603420|1604411 overexpression strains under vegetative and stressful conditions. (A) Quantitative RT-PCR (qRT-PCR) analysis of circRNA expression levels in WT strains of *T. marneffei* and four independent ectopic overexpression (OE) mutant strains (CO1–CO4). Data represent mean ± SD. (F) Phenotypic analysis of pm1_01:1603420|1604411 OE strains. Left panels show normal vegetative growth on SDA at 37°C. Right panels show representative spot assays and quantitative analysis under SDS medium at 37°C. Quantitative data represent the mean of three biological replicates. Error bars indicate SD.

In contrast, MSTRG.4967 overexpression mutants displayed a divergent phenotype, characterized by a marked sensitivity to Menadione at 25°C (Figures 8E–F) and a subtler yet detectable sensitivity at 37°C (Figures S7F). Given that MSTRG.4967 interacts with multiple modules involved in DNA repair, its overexpression may disrupt the homeostatic balance of these pathways, thereby compromising the cellular capacity to mitigate Menadione-induced reactive oxygen species (ROS) accumulation. Conversely, the overexpression of circRNA pm1_01:1603420|1604411 only marginally affected colony growth on SDS media (Figure S8). This mild phenotypic impact aligns with its modest network connectivity within the cell wall module (Figure 6A), suggesting a more specialized or redundant regulatory role under these specific stress conditions that is consistent with circRNAs predominantly exerting fine-tuning functions during the late phase of the dimorphic transition.

## Discussion

Since protein-coding variations and orphan gene repertoires are frequently insufficient to fully account for the complex adaptive traits of human fungal pathogens, unravelling how these organisms achieve diverse phenotypic plasticity points directly to regulatory network rewiring driven by non-coding elements^12^. To bridge this knowledge gap and systematically decode how upstream non-coding architectures govern fungal adaptation, we established a high-resolution temporal framework of non-coding RNA dynamics during reciprocal dimorphic transitions in *T. marneffei*, unveiling a multi-layered regulatory network wherein non-coding RNA transcripts govern specialized morphological transition and environmental adaptation, thereby offering a foundational paradigm for elucidating the molecular mechanisms driving fungal adaptational evolution and pathogenicity.

The temporal stratification of diverse non-coding RNA functions reveals a sophisticated division of regulatory labor driving transcriptomic rewiring during environmental adaptation, with lncRNAs predominating during early response phases and circRNAs governing later transition stages. While evolutionary investigations into non-coding transcripts remain relatively scarce, accumulating evidence indicates that conserved lncRNA transcripts often withstand stringent evolutionary selection, whereas circRNAs frequently exhibit pronounced species-specific importance^25,26^. Consistent with this paradigm, our observation of remarkable circRNA expression divergence despite underlying genomic sequence conservation in *Talaromyces*, where only a minor fraction of transcripts maintain conserved expression profiles, suggests that these molecules function as evolutionary capacitors facilitating rapid adaptation to thermal shifts, although distinguishing this adaptive function from non-functional transcriptional noise necessitates rigorous experimental validation. Furthermore, given that circRNA biogenesis relies on complex back-splicing mechanisms modulated by diverse RNA-binding proteins^24^, their delayed evolutionary emergence naturally positions them as potent sculptors of species-specific phenotypic traits. The selective enrichment of distinct RNA-binding protein motifs within their flanking sequences, which drives the generation of lineage-specific circular RNAs^14^, provides initial mechanistic insights into their evolutionary divergence, although these sequence-based predictions currently rely on mammalian-centric references and require direct empirical validation within fungal systems to firmly establish their contribution to evolutionary plasticity^24,27^.

The distinct phenotypic outcomes observed between hub lncRNA and circRNA perturbation experiments also highlight the profound evolutionary divergence in their regulatory functions. While the pronounced stress resistance conferred by the lncRNA overexpression of MSTRG.611 aligns with its architecture as a master network hub that integrates environmental sensing with core metabolic effectors during early phases, and while lncRNA MSTRG.4967 uniquely modulates oxidative stress sensitivity, the subtle phenotypic shifts associated with the circRNA pm1_01:1603420|1604411 reflect a fundamentally different mode of action. Specifically, circRNAs operate as late-acting modulators executing fine-tuned regulatory adjustments during morphological conversion, meaning their experimental perturbation yields nuanced rather than sweeping outcomes. This contrast illustrates a tiered evolutionary division of labor where lncRNA drive robust, broad spectrum stress preparation, whereas circRNAs provide precise, lineage-specific refinement. Although genetic disruption remains challenging due to technical barriers in non-model fungi such as *T. marneffei*, this systems-level dichotomy positions the non-coding genome as a multi-layered capacitor for transcriptional plasticity, establishing distinct intervention paradigms wherein early-acting lncRNAs serve as primary targets for broad-spectrum attenuation, while late-acting circRNAs offer specialized candidates for lineage-specific precision control against specialized pathogens.

Furthermore, moving beyond transcribed non-coding RNAs, non-coding genomic regions play a foundational pivotal role in driving evolutionary divergence^12^. The targeted enrichment of C6 zinc cluster transcription factor binding motifs exclusively within upregulated genes of *T. marneffei* elucidates the mechanistic basis of this regulatory divergence. The coordinated activation of these elements points to the emergence of a specialized regulatory module that couples thermal sensing with downstream metabolic reprogramming. From an evolutionary perspective, the notable absence of this conserved signature in phylogenetically proximate species suggests that this module arose via recent evolutionary innovations driven predominantly by *cis*-regulatory sequence divergence rather than novel transcription factor acquisition. This non-coding sequence evolution exemplifies how lineage-specific rewiring of promoter architectures provides a flexible evolutionary mechanism for acquiring complex thermal adaptations without requiring structural gene repertoire expansions^28–31^.

Future methodological horizons in this field involve several promising avenues of refinement. While Gaussian graphical model network analyses establish robust foundational frameworks, subsequent validation through alternative perturbation strategies will further enrich these regulatory models. Although targeted gene disruption of these central transcripts proved unsuccessful due to technical barriers intrinsic to non-coding genetic manipulation in non-model fungi *T. marneffei*, functional characterization successfully achieved overexpression of selected lncRNA and circRNA hubs, establishing a solid foundation for future systematic mutagenesis as genetic toolkits continue to evolve. Furthermore, broadening comparative temporal frameworks across diverse phylogenetic relatives will help capture additional response kinetics. Moving forward, prioritizing reporter assays for candidate *cis*-regulatory elements alongside advancing genetic tools will systematically examine hub lncRNAs and circRNAs, thereby clarifying their definitive roles in thermal adaptation and pathogenicity.

Collectively, our findings support a model wherein dimorphism of *T. marneffei* emerges through the evolutionary integration of multiple regulatory layers, encompassing *cis*-acting promoter architecture divergence, *trans*-acting lncRNA and circRNA regulation, and downstream effectors. Unlike structurally constrained protein-coding genes, rapidly evolving non-coding transcripts provide a flexible upstream layer driving gene regulatory network rewiring. Acting as modular connectors and lineage-specific modulators, lncRNAs and circRNAs enable rapid transcriptional restructuring under thermal stress, serving as primary drivers of phenotypic innovation. This intricate architecture explains the evolutionary rarity of thermal dimorphism, which requires simultaneous coordination across distinct regulatory systems.

## Methods

### Strains and culture conditions

The wild type *T. marneffei* strain PM1, which was isolated from a patient with culture-documented talaromycosis in Hong Kong^5^, was used in this study. Strains (mutants and wild type) of *T. marneffei* were grown on Sabouraud Dextrose Agar (SDA, Oxoid, Cambridge, UK) at 25℃ for seven days for conidia collection. The wild type *T. stipitatus* was purchased from CGMCC (3.15693, China General Microbiological Culture Collection Center) and cultured on SDB under 25℃ for 3 d for RNA extraction. The wild type *T. amestolkiae* was purchased from CICC (CICC 2700, China Center of Industrial Culture Collection) and cultured on SDB under 28℃ for 3 d for RNA extraction.

### Dynamic dimorphic transition

After culturing in SDB at 25°C or 37°C for 3 d, strains were switched to another temperature (such as 25°C transposed to 37°C). Smear observation was performed at 0, 12, 24, 36, 48, 72, 84 and 108 h after the temperature transition.

### Heat stress test of T. amestolkiae and T. stipitatus

After culturing in SDB at 25°C (*T. stipitatus*) or 28°C (*T. amestolkiae*) for 3d, strains were switched to 37°C. Smear observation and RNA extraction was performed at 0, 1 and 3 h after the temperature transition.

### rRNA depletion RNA-seq library preparation

The rRNA depletion RNA-seq was performed as previously described^14,19^. Briefly, rRNA genes in the *T. marneffei* genome were predicted using RNAmmer software. The rRNA sequences were segmented into consecutive 80-nucleotide probe sequences, with terminal regions shorter than 80 nucleotides adjusted to 60 nucleotides. All probes were synthesized by Qingke Biotechnology Co., Ltd. The synthesized rRNA-specific probes were subsequently mixed in equimolar ratios (18S/28S probes at a final concentration of 1 μM and 5S/5.8S probes at 0.1 μM). One microliter of probe mixture was combined with 2 μg total RNA and 2 μL hybridization buffer (750 mM Tris-HCl and 750 mM NaCl, pH adjusted to 8.0), with the total volume adjusted to 15 μL. The mixture was incubated at 95°C for 2 minutes, followed by gradual cooling to 22°C at a rate of -0.1°C/s. To the hybridization system, 1 μL thermostable RNase H (M0523, NEB) and 2 μL 10×buffer were added, adjusting the volume to 20 μL, and the mixture was incubated at 50°C for 30 minutes. Subsequently, 2.5 μL DNase I (M0303S, NEB) and 3 μL10× buffer were added to the digestion system, the volume was adjusted to 30 μL, and the mixture was incubated at 37°C for 30 minutes. For RNA purification, 75 μL of RNA magnetic beads (N412-01, Vazyme) were added to the depletion system, and RNA was eluted with 9 μL nuclease-free water. Strand-specific library construction was performed using the VAHTS Universal V8 RNA-seq Library Prep Kit (NR605-01, Vazyme) according to the manufacturer’s instructions, with fragmentation conditions set at 94°C for 5 minutes.

### Gene annotation of *Talaromyces*

Initially, nr and UniRef90 protein sequence databases were collected as reference datasets^32^. All open reading frames (ORFs) of predicted genes and transcripts were identified using TransDecoder software, followed by alignment of predicted ORFs against the nr and UniRef90 protein sequence databases using BLAST software^33^. After filtering ORFs with low alignment quality (E-value < 1×10⁻¹⁰), TransDecoder was employed to re-predict ORFs and map them to the genome to obtain predicted protein sequences. The predicted *T. marneffei*, *T. amestolkiae*, and *T. stipitatus* protein sequences were subsequently aligned against the nr protein sequence database, with low-quality alignment results (E-value < 1×10⁻¹⁰) filtered out. InterPro (IPR) domains of the *T. marneffei* gene sequences were predicted using InterProScan software, and the protein sequences were aligned against 14 databases including CDD, Coils, Gene3D, Hamap, MobiDBLite, PIRSF, PRINTS, Pfam, ProSitePatterns, ProSiteProfiles, SFLD, SMART, SUPERFAMILY, and TIGRFAM^34^. Finally, gene function prediction was performed by integrating BLASTp and InterProScan alignment results with the Gene Ontology (GO) database using Blast2GO software.

### RNA-seq data process

Raw Illumina RNA-seq data were subjected to sequencing quality assessment using MultiQC software^35^. Adapter removal and low-quality read filtering were performed using Trimmomatic^36^ with the following parameters: “ILLUMINACLIP:TruSeq3-PE-2.fa:2:30:10 LEADING:3 MINLEN:75 SLIDINGWINDOW:4:15”. The processed RNA-seq data were subsequently re-evaluated using MultiQC software to confirm data integrity and absence of anomalies. A genome-wide index was constructed using HISAT2 software, and quality-controlled RNA-seq reads were aligned to this index^37^. Strand-specific alignment files were generated by specifying the library type with the parameter --rna-strandness RF. Transcript assembly was performed using StringTie2 software with the --rf parameter to maintain strand specificity, resulting in gene annotation files (GTF format) containing comprehensive transcript information^38^. Differentially expressed genes (DEGs) were identified using EBSeq-HMM R package, with a corrected false discovery rate (FDR) < 0.05 and |log2(fold change) | ≥ 1 as thresholds^23^. Functional enrichment analyses were performed using the ClusterProfiler R package using aforementioned GO annotations^39^. Network of while transcriptomes was performed by GeneNet R packages using default parameters^40^.

### lncRNA identification and quantification

Gene annotation files were merged using TACO software^41^. The assembled transcripts were compared with the PM1 strain reference gene annotation file using GffCompare software, and transcripts with comparison codes “opxui” were retained as potential lncRNA candidates^42^. The coding potential of putative lncRNA transcripts was assessed using four independent tools: CPC2, CNCI, PfamScan, and FEELnc as previously descritped^19^. To minimize the influence of coding genes, only transcripts consistently identified as non-coding by all four software packages were retained as high-confidence lncRNAs as previously descripted^43^. GTF files from all samples were then merged using the merge module of StringTie2 software. Transcript quantification was performed using StringTie2 with the -e parameter to quantify transcripts present in the provided GTF file, generating gene and transcript expression matrices for each sample.

### circRNA identification and quantification

A reference genome index was constructed using BWA software^44^. Quality-controlled RNA-seq reads were subsequently aligned to the index using the mem module of BWA software with the parameter -T 19 to generate alignment files. Back-splice junction sites were identified from alignment files using CIRI2 software under the guidance of reference genome sequences and gene annotation files^45^. CircRNA identification was performed with the -high parameter setting, and the identified circRNAs were subsequently quantified using CIRIquant software^46^.

### Phylogenetic Tree Construction and Orthologous Gene Identification for Talaromyces Species

Genome sequences and annotation files for eight *Talaromyces* species including *T. amestolkiae* (ASM189636v1), *T. atroroseus* (ASM190759v1), *T. cellulolyticus* (ASM157146v2), *T. islandicus* (PIS), *T. stipitatus* (JCVI-TSTA1-3.0), *T. verruculosus* (ASM130527v1), *T. proteolyticus* (Talpro1), *T. rugulosus* (ASM1336875v1), with complete genome assemblies and gene annotations, along with Aspergillus fumigatus (ASM265v1) as an outgroup, were downloaded from NCBI. Single-copy orthologous genes were identified for *T. marneffei* and the aforementioned species using BUSCO software^47^. The obtained orthologous genes were subjected to multiple sequence alignment using MUSCLE software^48^, followed by alignment trimming with trimAl software to remove low-quality alignment positions^49^. Third, the multiple sequence alignment results were used as input for RAxML software^50^, employing the raxmlHPC-PTHREADS-AVX module with the ME algorithm for phylogenetic reconstruction, and an ultrametric tree was generated using the R package APE^51^. Finally, orthologous gene identification for *Talaromyces* species was performed using OrthoFinder software under the guidance of the phylogenetic tree to obtain the *Talaromyces* orthologous gene set^52^.

### Analysis of promoters of *Talaromyces*

Promoter sequences were defined as 2,000 bp regions upstream of gene annotations. When same-strand genes were present within this region, sequences were truncated to span from the end of the preceding gene to the start of the target gene. Only promoters exceeding 100 bp in length were included in subsequent analyses. Motif discovery was performed using MEME suite to identify enriched sequence motifs in promoters of upregulated, downregulated, and non-differentially expressed genes for each species separately^53^. The identified motifs were then compared against known fungal transcription factor binding sites in the JASPAR database using TOMTOM^53,54^. For regulatory element classification, promoter sequences from all three gene expression categories (upregulated, downregulated, and non-differentially expressed) were merged into a unified dataset. Differential expression patterns were used as class labels, with upregulated genes designated as potential enhancer-containing promoters, downregulated genes as silencer-containing promoters, and non-differentially expressed genes as neutral regulatory regions. A Random Forest classifier was trained on sequence features including nucleotide composition, dinucleotide frequencies, CpG content, GC content, tandem repeat scores, and conserved motif presence to predict regulatory element types. Model performance was evaluated using cross-validation and classification accuracy metrics.

### Conservation analysis of circRNAs

Since the full-length sequences of circRNAs are nearly identical to those of their cognate linear genes except for the back-splicing junction sequences, we focused on the regions spanning ±20 nucleotides flanking the back-splicing junctions of each circRNA. Sequences spanning 20 bp upstream and downstream of BSJs in the *T. marneffei* genome were extracted and aligned against the other eight Talaromyces genomes using the blastn module of BLAST software. Filtering was performed with the following thresholds: e-value <1×10⁻⁶, length ≥15, percent identity ≥80, identity count ≥12, mismatches ≤2, gaps ≤2, and upstream-downstream alignment distance ≤6,000 bp (95th percentile of *T. marneffei* gene length). CircRNA BSJs meeting these criteria were considered genomic-conserved BSJs. Intergenic circRNA sequences from *T. marneffei* were extracted and aligned against the other eight Talaromyces genomes using the blastn module of BLAST software, with filtering thresholds of e-value <1×10⁻⁶ and percent identity >60%. CircRNAs meeting these criteria were considered genomic-conserved intergenic circRNAs. Synteny analysis was performed on conserved circRNAs to identify upstream and downstream protein-coding genes flanking intergenic circRNAs. The conservation of genes flanking intergenic circRNAs was assessed and classified using orthologous gene results. CircRNAs with both ends conserved and detected in *T. amestolkiae* or *T. stipitatus* rRNA-depletion RNA-seq datasets were classified as transcriptomic-conserved circRNAs.

### Conservation analysis of lncRNAs

The full-length lncRNA sequences were subjected to BLAST analysis against genomes of all Talaromyces species, with filtering thresholds of e-value <1×10⁻^4^, percent identity > 50%, coverage > 30% and hit length >50 nt were considered genomic-conserved lncRNAs. Additionally, the full-length lncRNA sequences were compared via BLAST against lncRNAs newly identified in this study from *T. amestolkiae* and *T. stipitatus*, with filtering thresholds of e-value <1×10⁻^4^, percent identity > 50%, coverage > 30% and hit length >50 nt were considered as transcriptomic-conserved lncRNAs.

### Motif analysis of flanking sequences of circRNAs

To identify sequence motifs associated with back-splice junction (BSJ) formation, sequences spanning ±25 nt flanking each BSJ site were extracted. Motif discovery was performed using STREME software^53^, with sequences spanning ±25 nt around all exon boundaries serving as the background control set. Significantly enriched motifs were subsequently compared against established RNA-binding protein databases (CISBP-RNA, RNAcompete, and YEASTRACT+) using TOMTOM software^55–57^. Only motifs with *q*-values < 0.05 were retained as statistically significant findings, ensuring robust identification of potential regulatory elements involved in circRNA biogenesis.

### Overexpression of target ncRNAs

To achieve overexpression, the full-length sequences of lncRNA and circRNA (including flanking splicing sites for the latter) were amplified from genomic DNA and ligated into the pSilent-1 plasmid (Supplemental Table S2) following previously described protocols^14^. The integrity of the resulting constructs was verified via Sanger sequencing. The recombinant plasmids were then introduced into the spores via electroporation using a Bio-Rad system using the default program for yeast. Transformants were selected on SDA supplemented with hygromycin and subsequently validated by PCR analysis.

### Stress assays of mutants

The physiological responses of the mutants were evaluated under various stress conditions, including SDS (0.0005%–0.02% w/v), NaNO_2_ (0.5–8 mM), Congo Red (1–75 μM), KCl (0.3–1 M), sorbitol (0.5–2 M) and Menadione (10–150 μM). Spore suspensions were prepared at an initial concentration of 1×10^7^/μL. and subsequently subjected to a 10-fold serial dilution series. For each stress condition, 3 μL of each dilution was spotted onto the agar plates in ascending order of concentration. Each treatment was performed in triplicate. The cultures were incubated at 25°C and 37°C for 5 and 7 days, respectively, after which colony morphologies were photographed and colony diameters were quantified using Fiji software^58^.

### Data analysis and visualization

All data analyses were performed using R v4.1.2. Visualizations were performed using ggplot2 v3.3.6 except mentioned separately.

### Statistical tests

All statistical tests are completed in R v4.1.2, and the corresponding test methods and statistics are marked in the corresponding text.

## Acknowledgement

This work was funded by National Natural Science Foundation of China (32170091, Ence Yang and 32600273, Xueyan Hu).

## Author contributions

Ence Yang designed and supervised the project. Xueyan Hu performed bioinformatics analysis. Xueyan Hu and Juan Wang performed fungal experiments and molecular experiments. Xueyan Hu wrote the manuscript along with the revise of Ence Yang. All authors approved the final version of the manuscript.

## Availability of data and materials

The rRNA-depleted RNA-seq data in this study are deposited in the National Genomics Data Center BIG Sub system (https://ngdc.cncb.ac.cn/gsub/) under accession number PRJCA073567. Access to the data requires contacting the corresponding author.

## Additional information

### Supplemental Figures and Tables

**Table S1.** Genes of dimorphic-related pathways.

**Table S2.** Primers used in this study.

